# Reducing brain kynurenic acid synthesis precludes kynurenine-induced sleep disturbances

**DOI:** 10.1101/2022.11.21.517377

**Authors:** Katherine M. Rentschler, Snezana Milosavljevic, Annalisa M. Baratta, Courtney J. Wright, Maria V. Piroli, Zachary Tentor, Homayoun Valafar, Christian O’Reilly, Ana Pocivavsek

## Abstract

Patients with neurocognitive disorders often battle sleep disturbances. Kynurenic acid (KYNA) is a tryptophan metabolite of the kynurenine pathway implicated in the pathology of these illnesses. Modest increases in KYNA, an antagonist at glutamatergic and cholinergic receptors, result in cognitive impairments and sleep dysfunction. We explored the hypothesis that inhibition of the KYNA synthesizing enzyme, kynurenine aminotransferase II (KAT II), may alleviate sleep disturbances. At the start of the light phase, adult male and female Wistar rats received systemic injections of either i) vehicle, ii) kynurenine (100mg/kg; i.p.), iii) the KAT II inhibitor, PF-04859989 (30 mg/kg; s.c.), or iv) PF-04859989 and kynurenine in combination. Kynurenine and KYNA levels were evaluated in the plasma and brain. Separate animals were implanted with electroencephalogram (EEG) and electromyogram (EMG) telemetry devices to record polysomnography and evaluate vigilance states wake, rapid eye movement sleep (REMS) and non-REM sleep (NREMS) following each treatment. Kynurenine challenge increased brain KYNA and resulted in reduced REMS duration, NREMS delta power and sleep spindles. PF-04859989 reduced brain KYNA formation when given prior to kynurenine, prevented disturbances in REM sleep and sleep spindles, and enhanced NREM sleep. Our findings suggest that reducing KYNA in conditions where the kynurenine pathway is activated may serve as a potential strategy for improving sleep dynamics.

## Introduction

Healthy sleep promotes synaptic plasticity and is necessary for the maintenance of newly formed memories (Aton et al., 2009). Cortical electroencephalography (EEG) combined with electromyography (EMG) is readily employed to distinguish wake from sleep states. Non-rapid eye movement sleep (NREMS) presents as high amplitude cortical delta oscillations (0.5-4 Hz) from thalamocortical networks and rapid eye movement sleep (REMS) exhibits high amplitude frequency theta oscillations (4-8 Hz) from corticolimbic networks with muscle atonia. NREMS is largely implicated in procedural memory, and REMS sleep is considered critical to synaptic pruning, behavioral refinement, and memory consolidation (Abel, Havekes, Saletin, & Walker, 2013; Boyce, Williams, & Adamantidis, 2017). Patients with neurocognitive disorders often experience sleep dysregulation such as fragmented sleep, insomnia, poor quality of sleep, aberrant sleep spindles, REMS behavior disorders, and greater symptom severity following sleep disruption (Lucey et al., 2021; Pocivavsek & Rowland, 2018). Despite the integral relationship between sleep and cognition, there is currently a lack of FDA approved therapeutic options suitable to treat both cognitive symptoms and sleep disturbances.

To investigate a novel pharmacological intervention, we presently explore the impact of inhibiting kynurenine aminotransferase II (KAT II) on sleep physiology in rodents. KAT II is the primary synthesizing enzyme for brain-derived kynurenic acid (KYNA), a metabolite of the kynurenine pathway (KP) of tryptophan metabolism (see **Figure 1A**). The essential amino acid tryptophan is pivotal to circadian rhythmicity via the regulatory role of serotonin and melatonin in sleep and arousal (Rapoport & Beisel, 1968). However, only about 5% of tryptophan is degraded through the melatonin pathway. The vast majority of tryptophan is metabolized via the KP, which generates the small neuroactive molecule KYNA that we hypothesize serves a critical role in modulating sleep (Milosavljevic, Smith, Wright, Valafar, & Pocivavsek, 2023; Pocivavsek, Baratta, Mong, & Viechweg, 2017; Rentschler et al., 2021).

**Figure 1.**
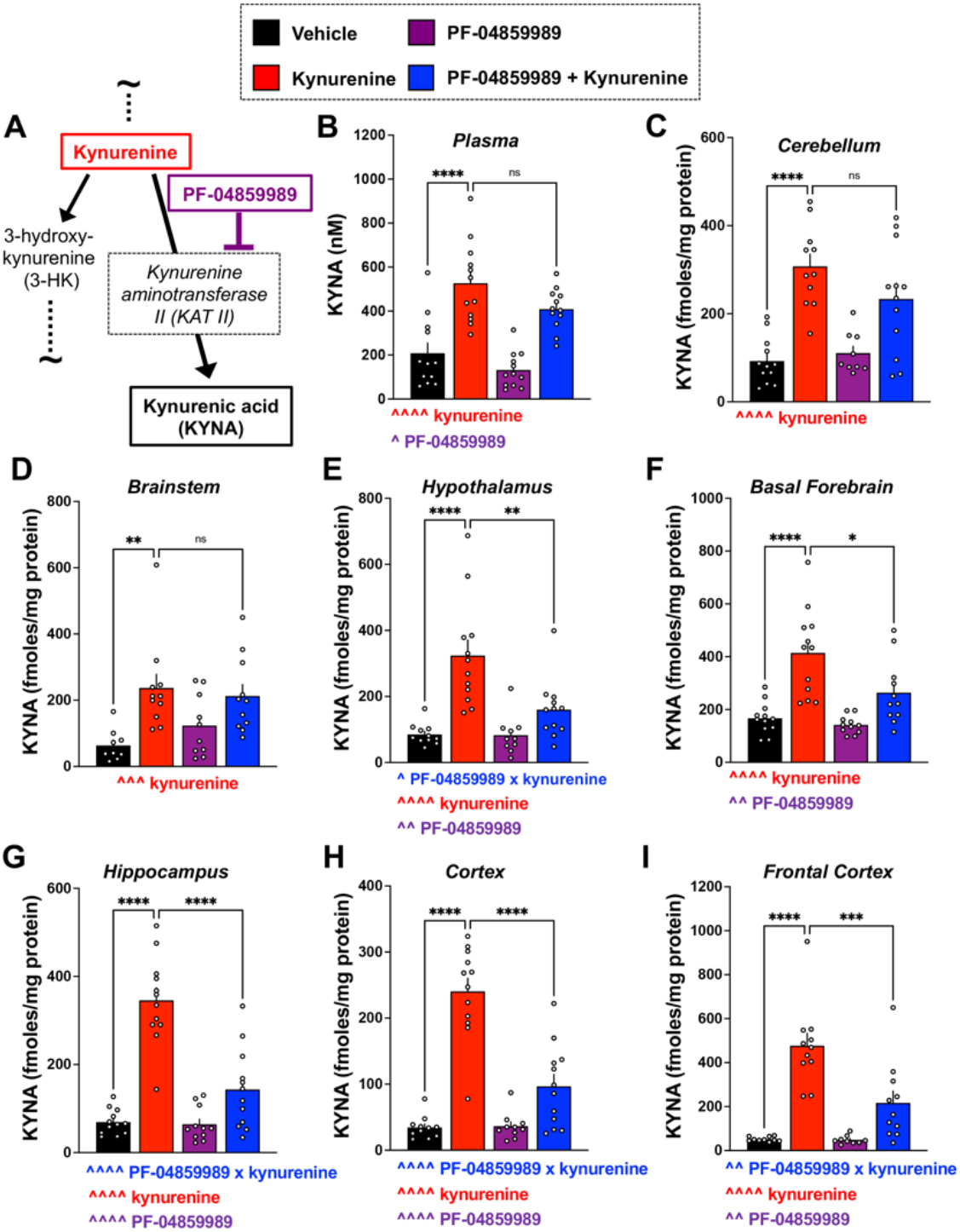
KYNA levels in plasma and across brain regions are increased after the acute kynurenine challenge, and reversed after pretreatment with the KAT II inhibitor, PF-04859989. (A) Schematic representation of truncated kynurenine pathway demonstrating key metabolites and pharmacological tools used in the present study. Adult rats were peripherally injected at Zeitgeber time (ZT) 0 with kynurenine (100 mg/kg) to induce *de novo* kynurenic acid (KYNA) formation. PF-04859989 (30 mg/kg), systemically active KAT II inhibitor, was given 30 minutes prior at ZT 23.5. Tissues were harvested at ZT 2. KYNA levels were evaluated in the (B) plasma; (C) cerebellum; (D) brainstem; (E) hypothalamus; (F) basal forebrain; (G) hippocampus; (H) cortex; (I) frontal cortex. All data are mean ± SEM. Two-way ANOVA analyses: ^P<0.05, ^^P<0.01, ^^^^P<0.0001; Bonferroni’s post hoc test: *P<0.05, **P<0.01, ***P<0.001, ****P<0.0001. N = 9-12 per group.

Elevations in KYNA disrupt cholinergic and glutamatergic-dependent processes, including learning and memory, and restoring a balance in KYNA promotes cognitive function (Kozak et al., 2014; Pocivavsek et al., 2011). Clinical and physiological significance for studying elevations in KYNA stems from growing evidence that increased KYNA is found in the brains of individuals with systemic infections, neurocognitive disease, and psychiatric disorders (Linderholm et al., 2012; Miller, Llenos, Dulay, & Weis, 2006; Sathyasaikumar et al., 2011; Schwarcz, Bruno, Muchowski, & Wu, 2012). To combat cognitive dysfunction associated with these illnesses, translational studies have evaluated pharmacological approaches to reduce KYNA in the brain by inhibiting the synthesizing enzyme KAT II (Kozak et al., 2014; Pellicciari et al., 2006; Pocivavsek & Erhardt, 2023; Pocivavsek et al., 2011).

Our present goal was to pharmacologically inhibit KAT II as a mechanistic strategy to i) reduce *de novo* KYNA production after acute treatment with kynurenine, the direct bioprecursor of KYNA, and ii) overcome sleep disturbances induced by excessive KYNA (Pocivavsek et al., 2017). We selected PF-04859989, a brain penetrable and irreversible KAT II inhibitor (Klausing, Fukuwatari, Bucci, & Schwarcz, 2020; Kozak et al., 2014) and confirmed its effectiveness to reduce *de novo* KYNA formation when administered prior to kynurenine challenge. We thereby hypothesize that KAT II inhibition with PF-04859989 will effectively attenuate dysregulated sleep-wake behavior in response to an acute kynurenine challenge.

## METHODS

### Animals

Adult male (250-350 g) and female (200-300 g) Wistar rats were obtained from Charles River Laboratories. Animals were kept in a facility fully accredited by the American Association for the Accreditation of Laboratory Animal Care on a 12/12 h light-dark cycle, where lights on corresponded to zeitgeber time (ZT) 0 and lights off to ZT 12. All protocols were approved by Institutional Animal Care and Use Committees and were in accordance with the National Institutes of Health *Guide for the Care and Use of Laboratory Animals*.

### Chemicals

L-kynurenine sulfate (“kynurenine”, purity 99.4%) was obtained from Sai Advantium (Hyderbad, India). PF-04859989 was obtained from WuXi AppTec (purity 95%; Shanghai, China). Other chemicals were obtained from various suppliers and were of the highest commercially available purity. For intraperitoneal (i.p.) injections, kynurenine was prepared daily by dissolving in phospho-buffered saline and adjusting the pH to 7.4 with 0.1N NaOH. For subcutaneous (s.c.) injections, PF-04859989 was prepared daily by dissolving in ultrapure water.

### Biochemical Analysis

Male and female rats were treated acutely with vehicle (water, s.c.) or PF-04859989 (30mg/kg, s.c.) at ZT 23.5 and vehicle (saline, i.p.) or kynurenine (100 mg/kg, i.p.) at ZT 0. Animals were euthanized at ZT 2 by CO_2_ asphyxiation. Whole trunk blood was collected in tubes containing 25 μl K3-EDTA (0.15%) as an anticoagulant, and centrifuged (1000 x g, 10 min) to separate plasma. Rapidly removed brains were put on ice, and the following regions were dissected: cerebellum, brainstem, hypothalamus, basal forebrain, hippocampus, cortex, and frontal cortex. All samples were promptly snap-frozen on dry ice and stored at -80°C until biochemical analyses. Kynurenine and KYNA levels were determined by high-performance liquid chromatography (HPLC) (Wright et al. 2021; Buck et al., 2020). Briefly, plasma samples were diluted (1:10) in ultrapure water, and tissue samples were weighed and sonicated (1:5, w/v) in ultrapure water before being further diluted (final concentration: 1:10, w/v). One hundred μl of sample, either diluted plasma or tissue homogenate, was acidified with 25 μl of 6% (plasma) or 25% (tissue) perchloric acid, centrifuged (12000 x g, 10 min), and 20 μl of the supernatant was subjected to HPLC. Kynurenine and KYNA were isocratically eluted from a ReproSil-Pur C18 column (4 x 100 mm; Dr. Maisch GmbH, Ammerbuch, Germany) using a mobile phase containing 50 mM sodium acetate and 5% acetonitrile (pH adjusted to 6.2 with glacial acetic acid) at a flow rate of 0.5 ml/min. Zinc acetate (500 mM) was delivered after the column at a flow rate of 0.1 ml/min. Kynurenine (excitation: 365 nm, emission: 480 nm, retention time ∼7 minutes) and KYNA (excitation: 344 nm, emission: 398 nm, retention time ∼13 minutes) were detected fluorometrically (Waters Alliance, 2475 fluorescence detector, Bedford, MA) in the eluate. Data were integrated and analyzed using Empower 3 software (Waters). The total concentration of protein in tissue samples was determined by the Lowry protein assay .

### Surgery

Under isoflurane anesthesia (5% induction, 2-3% maintenance), animals were placed in a stereotaxic frame (Stoelting, Wood Dale, IL) for the surgical implantation of EEG/EMG telemetry transmitters (PhysiolTel HD-S02; Data Science International, St. Paul, MN) according to previously published protocols (Pocivavsek et al., 2017; Rentschler et al., 2021). The transmitter was inserted intraperitoneally through a dorsal incision of the abdominal region. An incision at the midline of the head was made to anchor two EEG leads to their corresponding surgical screws (Plastics One, Roanoke, VA) placed into 0.5 mm burr holes at 2.0 mm anterior/+1.5 mm lateral and 7.0 mm posterior/−1.5 mm lateral relative to bregma and secured with dental cement. Two EMG leads were inserted and sutured into the dorsal cervical neck muscle, separated by 1.0 mm. The skin on top of the head was sutured, the abdominal incision was closed with wound clips, and animals recovered for ≥ 7 days prior to experimentation.

### Vaginal Lavage and Estrous Cycle Tracking

To track the estrous cycle, lavage of the vagina in females was taken using saline-filled transfer pipettes. Samples were viewed with a SeBa digital microscope (LAXCO Inc., Mill Creek, WA, USA) within 30 minutes of sample collection. Best efforts were made to avoid drug treatments during proestrus, wherein sleep and wake parameters are vastly different from other cycle stages for female subjects (Swift et al., 2020).

### Drug Treatments for Sleep Data Acquisition

*Sleep Study #1:* Counterbalanced vehicle (saline, i.p.) or kynurenine (100 mg/kg, i.p.) injections were given 96 h apart at ZT0. Polysomnography data were acquired for 24 h after each treatment. *Sleep Study #2:* On the first day of the experiment, animals were injected with vehicle (water, s.c.) at ZT 23.5 and vehicle (saline, i.p.) at ZT 0 to obtain baseline sleep data (denoted “vehicle” in graphs). After 96 h, animals received PF-04859989 (30 mg/kg, s.c.) at ZT 23.5 and either vehicle (saline, i.p.; denoted “PF-04859989 + vehicle” in graphs) or kynurenine (100 mg/kg, i.p.; denoted “PF-04859989 + kynurenine” in graphs) at ZT 0. The experimental schedule was counterbalanced and repeated, such that each animal received PF-04859989 + vehicle and PF-04859989 + kynurenine injections. Sleep data acquisition occurred for 24 h after ZT 0 treatment. Treatments were spaced 96 h apart to record sleep within the same estrous phase for female subjects.

### Sleep Data Acquisition and Analyses

Animals undergoing sleep recordings were left undisturbed in a designated room. EEG/EMG data and home cage activity were recorded using Ponemah 6.10 software (DSI) with a continuous sampling rate of 500 Hz (Milosavljevic et al., 2023; Pocivavsek et al., 2017; Rentschler et al., 2021). All digitized signal data were scored offline with NeuroScore 3.0 (DSI) using 10-second epochs as wake (low-amplitude, high-frequency EEG combined with high-amplitude EMG), NREMS (high-amplitude, low-frequency EEG with low-amplitude EMG), or REMS (low-amplitude, high-frequency EEG with very low-amplitude EMG). **Supplemental Information** includes representative hypnograms and continuous EEG and EMG recordings. Transitions among vigilance states were scored when two or more 10-s epochs of the new state were noted. Parameters assessed included total vigilance state duration, number of bouts, average bout duration, number of transitions between vigilance states, bout length distribution and spectral power frequencies during REMS and NREMS. Discrete fast Fourier transform (DFT) was used to calculate the EEG power spectrum for each epoch in the delta (0.5 – 4 Hz), theta (4 – 8 Hz), alpha (8 – 12 Hz), sigma (12 – 16 Hz), and beta (16 – 20 Hz) frequency ranges and normalized to the total power per subject. Sleep spindles were automatically detected based on previously employed methodology (Uygun et al., 2019) and verified in NeuroScore by manual expert scoring detection (see **Supplemental Information**).

### Statistical Analyses

Normality of data was assessed with Shapiro-Wilk test and data were visually inspected using Q-Q plots to confirm a relative bell-shaped distribution and the absence of outliers. For biochemical analyses, comparisons were made using a two-way analysis of variance (ANOVA) with kynurenine and/or PF-04859989 treatment as a between-subject factors. Significant main effects were followed by Bonferroni’s correction with an adjusted alpha for multiple comparisons. To evaluate sex as a biological variable in sleep architecture data (vigilance state durations, bout number, and average bout duration), we initially employed three-way repeated measures (RM) ANOVA with treatment and ZT as within-subject factors and sex as a between-subject factor. As no main effects of sex were determined (see **Supplemental Table 1**), the remaining analyses were conducted with sexes combined. Durations were evaluated in 1-hr bins, followed by 6-hr bins, while bout number and average bout duration were evaluated in 6-hr bins employing two-way RM ANOVA with treatment and ZT as within-subject factors. REMS theta power spectra and NREMS delta power spectra were evaluated during the light phase by two-way RM ANOVA with frequency and treatment as within-subject factors. Spindle density, average spindle duration, and spindle amplitude were evaluated by two-way RM ANOVA with treatment and ZT as within-subject factors. Power spectra during spindles were evaluated with two-way RM ANOVA with frequency and treatment as within-subject factors. Vigilance state transitions were evaluated by paired Student’s t-test. Relative cage activity was analyzed by three-way RM ANOVA with treatment and ZT as within-subject factors, and sex as a between-subject factor. Post hoc analysis was conducted with Bonferroni’s correction in Sleep Study #1 and Dunnett test in Sleep Study #2, to compare drug treatments to vehicle treatment condition. Percent of vehicle calculations (REMS duration, NREMS slow-wave spectral power and spindle density) allowed comparisons between Sleep Study #1 and Sleep Study #2, thereby comparing PF-04859989 + kynurenine to kynurenine treatment alone with unpaired Student’s t-test. All statistical analyses were performed using Prism 9.0 (GraphPad Software, La Jolla, CA, USA) and significance was defined as P < 0.05.

## RESULTS

### Systemic administration of KAT II inhibitor PF-04859989 attenuates elevation in brain KYNA levels induced by acute kynurenine challenge

We first confirmed that acute systemic kynurenine treatment at ZT 0, start of the light phase, elevates plasma kynurenine levels at ZT 2 compared to vehicle treatment (**Supplemental Table 2**). Kynurenine-treated rats exhibited a 2.8-fold increase in plasma kynurenine levels compared with vehicle-treated rats (P<0.0001). As kynurenine promptly enters the brain from the blood, we confirmed significant elevation in brain kynurenine levels in the cerebellum (6.2-fold; P<0.0001), brainstem (2.3-fold; P<0.0001), hypothalamus (2.5-fold; P<0.0001), basal forebrain (3.9-fold; P<0.0001), hippocampus (5.2-fold; P<0.0001), cortex (4.1-fold; P<0.0001), and frontal cortex (3.6-fold; P<0.0001) compared to vehicle. Levels of kynurenine in plasma and brain remained unchanged after pretreatment with PF-04859989.

Upon kynurenine treatment at ZT 0, plasma KYNA increased 2.5-fold compared to vehicle-treated rats (*F*_1,43_ = 73.71, P<0.0001) (**Figure 1B**). Across brain regions (**Figure 1C-I**), we similarly determined that kynurenine treatment elevated KYNA content in the cerebellum (3.3-fold; *F*_1,40_ = 40.95, P<0.0001), brainstem (3.7-fold; *F*_1,37_ = 15.74, P<0.0001), hypothalamus (3.8-fold; *F*_1,41_ = 27.20, P<0.001), basal forebrain (2.5-fold; *F*_1,42_ = 32.66, P<0.0001), hippocampus (5.0-fold; *F*_1,43_ = 71.33, P<0.0001), cortex (7.0-fold; *F*_1,41_ = 80.26, P<0.0001), and frontal cortex (9.2-fold; *F*_1,37_ = 49.44, P<0.0001). PF-04859989 significantly impacted KYNA levels noted as a main effect of treatment peripherally in plasma (*F*_1,43_ = 4.848, P<0.05) and also in the hypothalamus (*F*_1,41_ = 7.497, P<0.01), basal forebrain (*F*_1,42_ = 7.396, P<0.01), hippocampus (*F*_1,43_ = 24.23, P<0.0001), cortex (*F*_1,41_ = 22.71, P<0.0001), and frontal cortex (*F*_1,37_ = 9.691, P<0.0001). Pretreatment with PF-04859989 attenuated the increase in KYNA following acute kynurenine challenge, noted by a PF-04859989 x kynurenine interaction in the hypothalamus (*F*_1,41_ = 7.076, P<0.05), hippocampus (*F*_1,43_ = 22.00, P<0.0001), cortex (*F*_1,41_ = 23.98, P<0.0001), and frontal cortex (*F*_1,37_ = 9.248, P<0.01). Post-hoc analysis noted a significant reduction in KYNA when PF-04859989 was given prior to kynurenine compared to kynurenine alone in the hypothalamus (P<0.01), basal forebrain (P<0.05), hippocampus (P<0.0001), cortex (P<0.0001), and frontal cortex (P<0.01). PF-04859989 pretreatment did not significantly reduce KYNA formation in the plasma, cerebellum and brainstem. In summary, **Table 1** indicates the percent increase in KYNA levels from vehicle with kynurenine treatment (top panel) and the percent decrease in KYNA levels when PF-04859989 was given prior to kynurenine challenge (bottom panel) in the plasma and individual brain regions investigated.

**Table 1.**
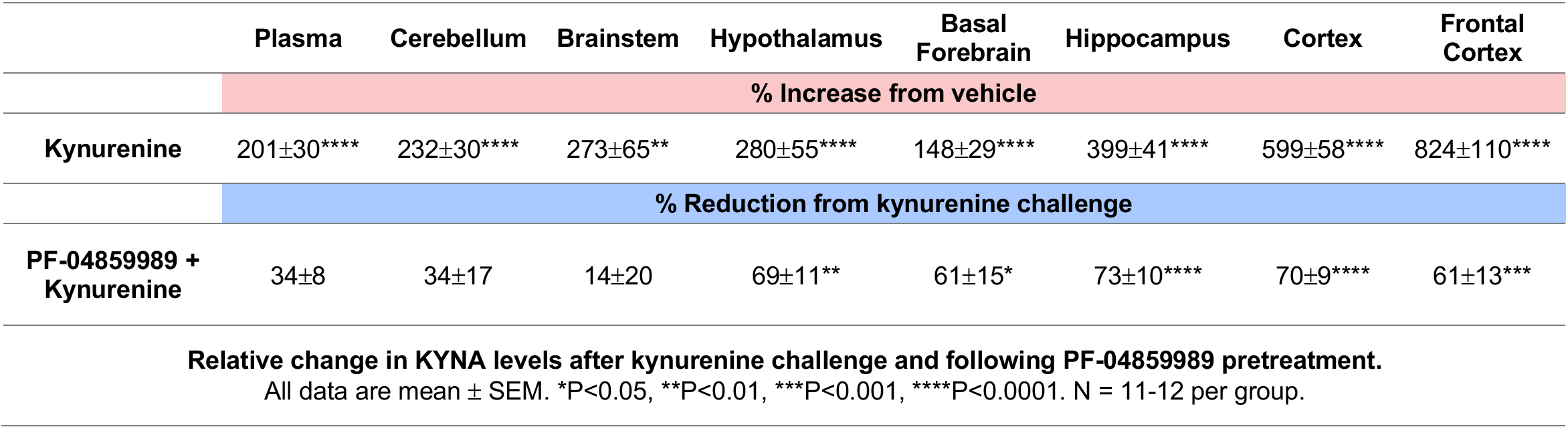
Relative change in KYNA levels after kynurenine challenge and following PF-04859989 pretreatment. Adult rats were peripherally injected at Zeitgeber time (ZT) 0 with kynurenine (100 mg/kg) to induce *de novo* kynurenic acid (KYNA) formation. PF-04859989 (30 mg/kg), systemically active KAT II inhibitor, was given 30 minutes prior at ZT 23.5. Tissues were harvested at ZT 2. All data are mean ± SEM. *P<0.05, **P<0.01, ***P<0.001, ****P<0.0001. N = 11-12 per group.

|  | Plasma | Cerebellum | Brainstem | Hypothalamus | Basal Forebrain | Hippocampus | Cortex | Frontal Cortex |
| --- | --- | --- | --- | --- | --- | --- | --- | --- |
|  | % Increase from vehicle |  |  |  |  |  |  |  |
| Kynurenine | 201±30**** | 232±30**** | 273±65** | 280±55**** | 148±29**** | 399±41**** | 599±58**** | 824±110**** |
|  | % Reduction from kynurenine challenge |  |  |  |  |  |  |  |
| PF-04859989 + Kynurenine | 34±8 | 34±17 | 14±20 | 69±11** | 61±15* | 73±10**** | 70±9**** | 61±13*** |
| Relative change in KYNA levels after kynurenine challenge and following PF-04859989 pretreatment.<br>All data are mean ± SEM. *P<0.05, **P<0.01, ***P<0.001, ****P<0.0001. N = 11-12 per group. |  |  |  |  |  |  |  |  |

### Kynurenine treatment reduces sleep and alters architecture, spectral power, and spindle characteristics

Sleep Study #1 was conducted in a cohort of rats used exclusively for sleep monitoring experiments. No main effects of sex were found when sex was included in the ANOVA of sleep data (**Supplemental Table 1)**. Vigilance state durations in 1-hr bins are shown separately for males and females in **Supplemental Figure 1**. For the remaining analysis of vigilance states, we combined sexes. When analyzed with 1-hr bins, REMS duration decreased significantly for kynurenine treatment was compared to vehicle treatment (ZT x treatment: *F*_23,437_ = 1.635, P<0.05; treatment: *F*_1,19_ = 15.14, P<0.01). The starkest REMS duration decrease occurred in the later portion of the light phase at ZT 10 (P<0.01)(**Figure 2A**). Analyses in 6-hr time bins supported the notion that REMS was disrupted with kynurenine challenge, particularly in the latter half of the light phase (ZT 6-12). REMS duration (treatment: *F*_1,19_ = 15.15, P<0.001)(**Figure 2B**) and REMS bouts (treatment: *F*_1,19_ = 8.510, P<0.01)(**Figure 2C**) were significantly reduced with kynurenine challenge. Parameters that were not significantly altered with kynurenine treatment related to REMS included the average duration of each REMS bout (**Supplemental Figure 2A**) and spectra power in the theta range during the light phase (**Supplemental Figure 2B**).

**Figure 2.**
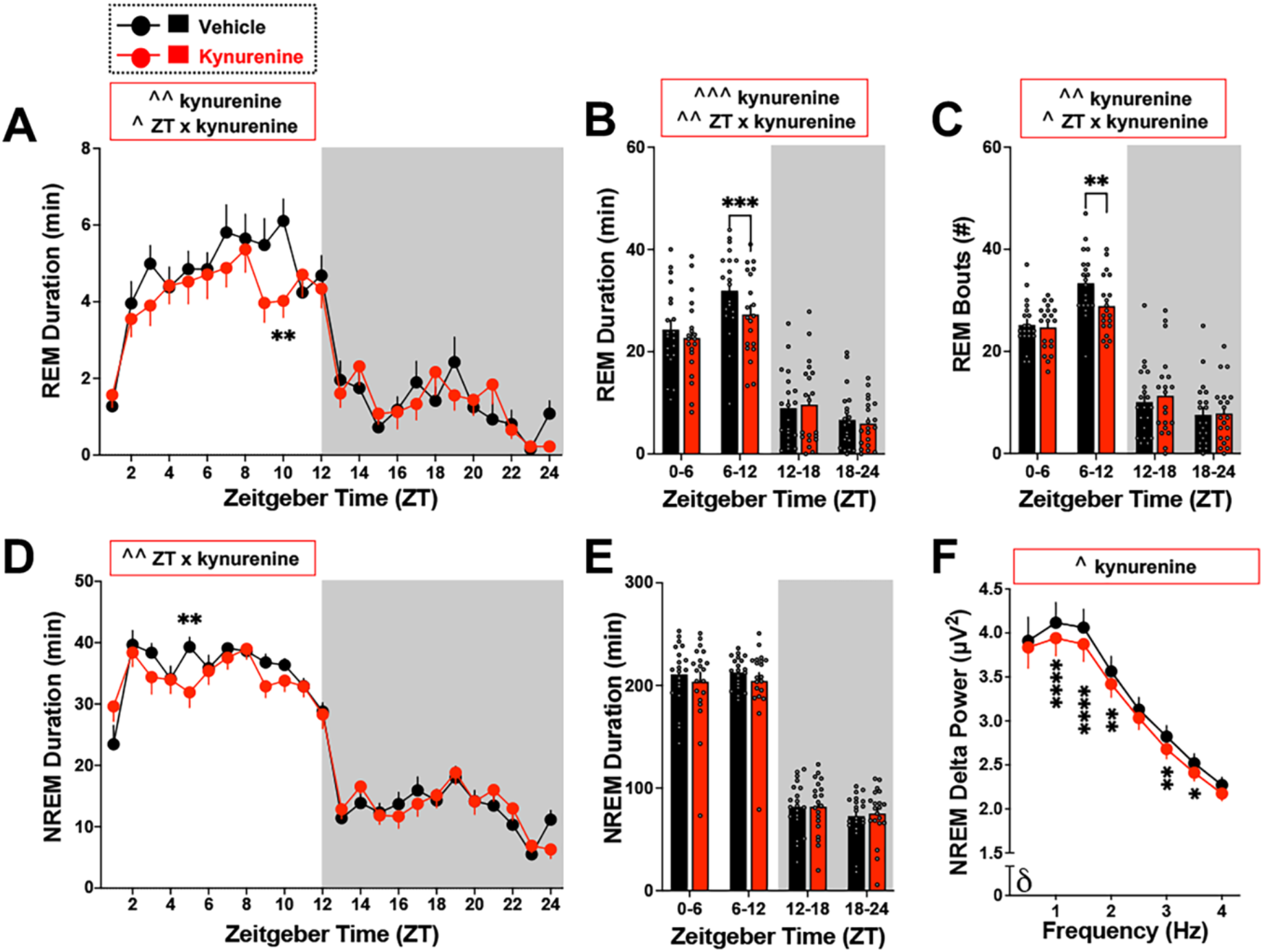
Kynurenine challenge reduces sleep and alters sleep architecture. Adult rats were treated with vehicle or kynurenine (100 mg/kg) at Zeitgeber time (ZT) 0. Data are mean ± SEM, analyzed by two-way RM ANOVA, with significance shown in graphs as ^P<0.05, ^^P<0.01, ^^^P<0.001 and Bonferroni’s post hoc test **P<0.01, ***P<0.001, ****P<0.0001. **(A)** 1-hr bins of REMS duration. ZT x treatment: *F*_23,437_ = 1.635, P<0.05; treatment: *F*_1,19_ = 15.14, P<0.01; ZT: *F*_23,437_ = 30.82, P<0.0001. **(B)** 6-hr bins of REMS duration. ZT x treatment: *F*_3,57_ = 4.162, P<0.01; treatment: *F*_1,19_ = 15.15, P<0.001; ZT: *F*_3,57_ = 99.50, P<0.0001. **(C)** 6-hr bins of REMS bout number. ZT x treatment: *F*_3,57_ = 3.74, P<0.05; treatment: *F*_1,19_ = 8.510, P<0.01; ZT: *F*_3,57_ = 112.9, P<0.0001. **(D)** 1-hr bins of NREMS duration. ZT x treatment: *F*_23,437_ = 2.032, P<0.01; ZT: *F*_23,437_ = 69.02, P<0.0001 **(E)** 6-hr bins of NREMS duration. ZT: *F*_3,57_= 247.3, P<0.0001 **(F)** NREMS delta spectral power during the entire light phase. Treatment: *F*_1,17_ = 4.615, P<0.05; Frequency: *F*_7,119_ = 64.15, P<0.0001. N = 20 per group (8 males, 12 females).

Kynurenine treatment induced changes to NREMS duration that varied across 1-hr bin analysis (treatment x ZT interaction: *F*_23,437_= 2.032, P<0.01). NREMS duration decreased significantly at ZT 5 (P<0.01) with kynurenine treatment compared to vehicle (**Figure 2D**). Analyses in 6-hr time bins revealed no significant changes in NREMS duration (**Figure 2E**), NREMS bout number (**Supplemental Figure 2C**), and average duration of NREMS bouts (**Supplemental Figure 2D**). Delta (0.5 – 4 Hz) spectral power was highest compared with theta, alpha, sigma, and beta during NREMS (light phase: *F*_38,722_ = 414.2, P<0.0001; data not shown). During the entire light phase, kynurenine treatment significantly reduced NREMS delta spectral power (treatment: *F*_1,17_ = 4.615, P<0.05; post-hoc 1 Hz: P<0.001; 1.5 Hz: P<0.0001; 2 Hz: P<0.01; 3 Hz: P<0.01; 3.5 Hz: P<0.05)(**Figure 2F**). Power spectra across 1-hr bin analysis is shown in Supplemental Figure 2E.

NREMS sleep spindles were detected during the light phase. Kynurenine treatment modified spindle density across 4-hr bin analysis (treatment x ZT interaction: *F*_2,16_= 7.648, P<0.01), with reduced spindle density from ZT0-4 (P<0.001)(**Figure 3A**). Average duration of each spindle (**Figure 3B**), peak to peak amplitude (**Figure 3C**) and the frequency distribution of spindle power (**Figure 3D**) remained unchanged when comparing kynurenine and vehicle treatments.

**Figure 3.**
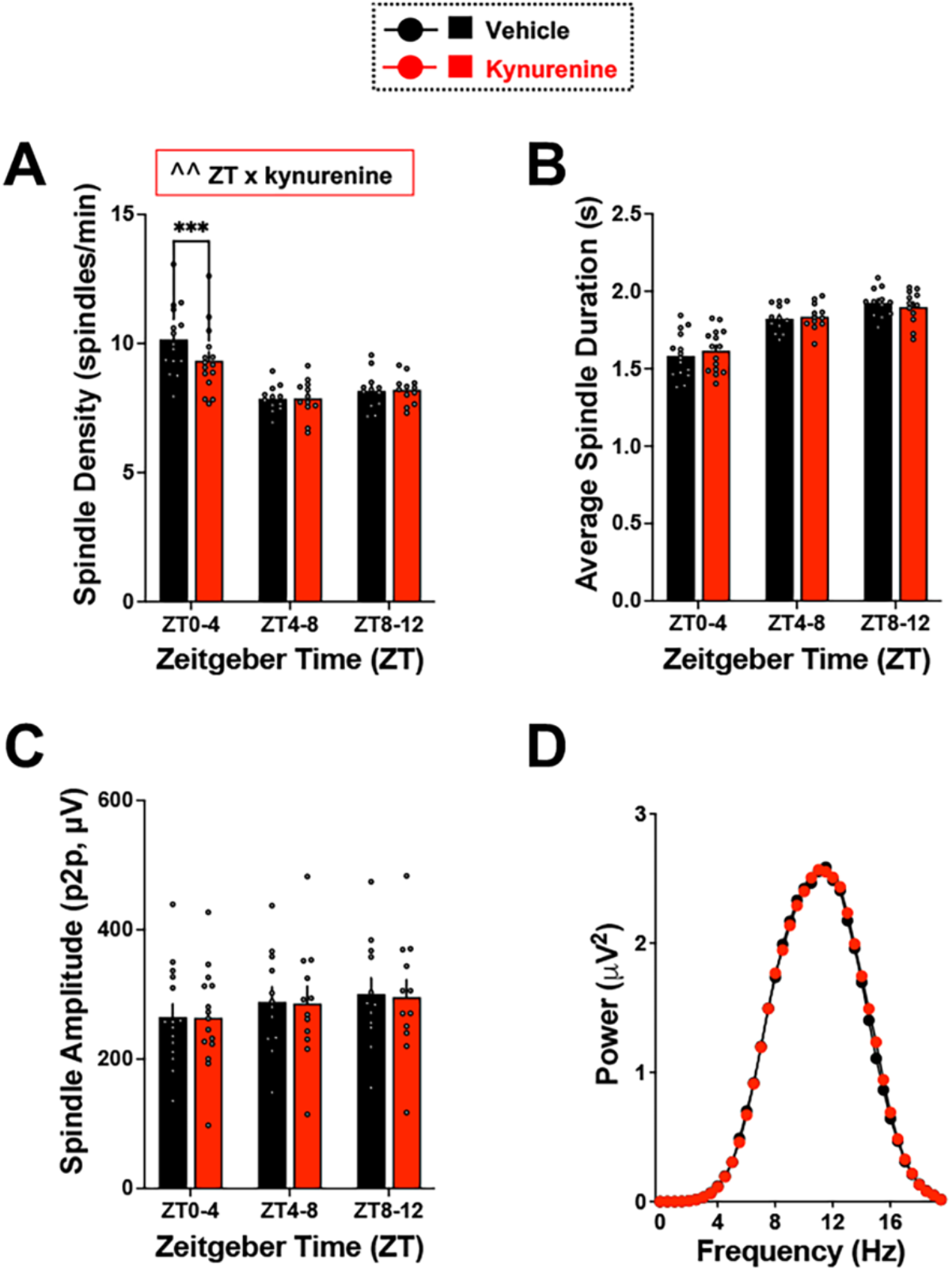
Kynurenine challenge reduces NREM sleep spindle density. Adult rats were treated with vehicle or kynurenine (100 mg/kg) at Zeitgeber time (ZT) 0. Sleep spindles were evaluated during the light phase. Data are mean ± SEM, analyzed by two-way RM ANOVA **(A)** NREMS spindle density (spindles/min NREMS). Treatment x ZT interaction: *F*_2,16_= 7.648, P<0.01. ZT: *F*_2,28_= 23.43, P<0.0001. **(B)** Average spindle duration. ZT: *F*_2,28_= 133.0, P<0.0001. **(C)** Spindle peak to peak (p2p) amplitude. ZT: *F*_2,28_= 4.023, P<0.05 **(D)** NREMS spindle power. Frequency: *F*_39,741_ = 185.0, P<0.0001. N = 20 per group (8 males, 12 females).

### Kynurenine challenge alters wake architecture

To understand how the observed changes in REMS and NREMS impacted arousal, wake state architecture and general home cage activity were evaluated. Analysis of wake durations in 1-hr bins revealed a kynurenine x ZT interaction (*F*_123,437_ = 2.078, P<0.01) such that kynurenine initially reduced wake duration within the first hour (P<0.01)**(Supplemental Figure 3A)**. When wake parameters were evaluated in 6-hr bins, we did not find any significant changes in wake duration (**Supplemental Figure 3B**), number of wake bouts (**Supplemental Figure 3C**), or average duration of each wake bouts (**Supplemental Figure 3D**). Of note, sex significantly impacted home cage activity (three-way ANOVA, *F*_1,347_ = 5.118, P<0.05), along with time of day (*F*_22,396_ = 25.40, P<0.0001), yet kynurenine treatment did not change home cage activity across time (**Supplemental Figure 3E**).

### Vigilance state transitions are impacted by kynurenine challenge

The stability of maintaining a wake or sleep episode upon kynurenine challenge was assessed by examining the number of transitions into and out of wake, NREMS sleep, and REMS sleep by light or dark phase (**Supplemental Table 3**). Kynurenine challenge reduced REMS to NREMS (P < 0.0001) and NREMS to REMS (P < 0.01) transitions during the light phase. In the subsequent dark phase, kynurenine treatment resulted in enhanced wake to NREMS (P < 0.05) and REMS to wake (P <0.05) transitions. Analysis of bout length distribution for the light phase showed that kynurenine treatment significantly decreased short NREMS bouts (<0.5 min) and REMS bouts (0.5-1 min)(**Supplemental Figure 4**).

### KAT II inhibitor PF-04859989 attenuates changes in sleep architecture and spindle dynamics induced by acute kynurenine challenge

Pretreatment with PF-04859989 resulted in reduced *de novo* KYNA synthesis in the brain with systemic kynurenine challenge (**Figure 1**), thus we next explored if PF-04859989 would readily reverse the consequences of kynurenine challenge on sleep architecture. Sleep Study #2 was conducted in a new cohort of adult rats (N = 18), and each subject received the following treatments: i) two vehicle injections (ZT 23.5 s.c., ZT 0 i.p.), ii) PF-04859989 at ZT 23.5 + vehicle at ZT 0, iii) PF-04859989 at ZT 23.5 + kynurenine at ZT 0. Our analysis focused on vigilance state parameters that were altered with kynurenine challenge, as reported above. Thus, focusing initially on REMS, we found that PF-04859989 pretreatment prevented kynurenine-induced REMS deficits, thereby stabilizing REMS duration (1-hr bins: **Figure 4A**; 6-hr bins: **Figure 4B**), REMS bout number (**Figure 4C**), average duration of each REMS bout (**Supplemental Figure 5A**), and theta spectral power during REMS (**Supplemental Figure 5B**) to vehicle treatment levels. PF-04859989 pretreatment did not influence NREMS duration (1-hr bins: **Figure 4D**; 6-hr bins: **Figure 4E**), NREMS bout number (**Supplemental Figure 5C**), and average duration of each NREMS bout (**Supplemental Figure 5D**). Of note, PF-04859989 treatment, with or without kynurenine, did not alter NREMS delta power, in 1-hr bin (**Supplemental Figure 5E)** or entire light phase (**Figure 4F**) analysis.

**Figure 4.**
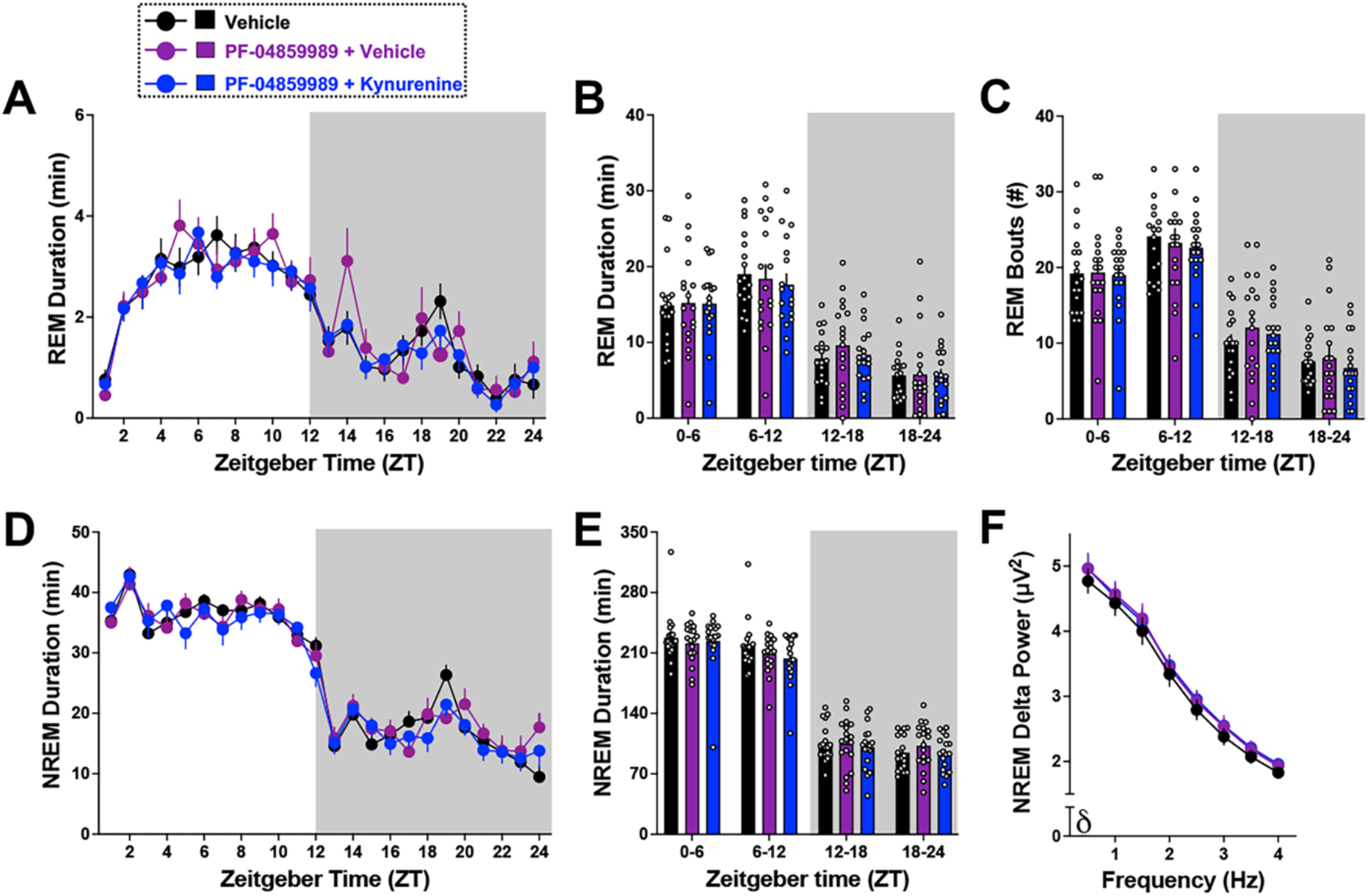
PF-04859989 treatment enhances NREM sleep architecture. Adult rats were treated with vehicle, PF-04859989 (30 mg/kg), or PF-04859989 + kynurenine (100 mg/kg) at the start of the light phase. Data are mean ± SEM, analyzed by two-way RM ANOVA, with significance shown in graphs as ^P<0.05, and Dunnett’s post hoc test *P<0.05 (PF-04859989 + vehicle vs. vehicle), ***P<0.001 (PF-04859989 + kynurenine vs. vehicle). **(A)** 1-hr bins of REMS duration. ZT: *F*_23,391_= 23.98, P<0.0001. **(B)** 6-hr bins of REMS duration. ZT: *F*_3,51_ = 62.44, P<0.0001. **(C)** 6-hr bins of REMS bout number. ZT: *F*_3,51_ = 76.02, P<0.0001. **(D)** 1-hr bins of NREMS duration. ZT: *F*_23,391_= 86.51, P<0.0001 **(E)** 6-hr bins of NREMS duration. ZT: *F*_3,51_= 304.4, P<0.0001 **(F)** NREMS delta spectral power during the entire light phase. Frequency: *F*_7,91_ = 135.0, P<0.0001. N = 18 per group (9 males, 9 females).

Evaluation of spindle dynamics revealed no significant differences in NREMS spindle density between treatments in Sleep Study #2 (**Figure 5A**), suggesting that PF-04859989 attenuated the reduction in NREMS spindle density elicited by kynurenine (comparison to **Figure 3A**). Average spindle duration and peak-to-peak amplitude were not impacted by PF-04859989 treatment (**Figure 5B-C**). Spectral power within the 10-15 Hz sigma range was also not impacted by PF-04859989 treatment (**Figure 5D**).

**Figure 5.**
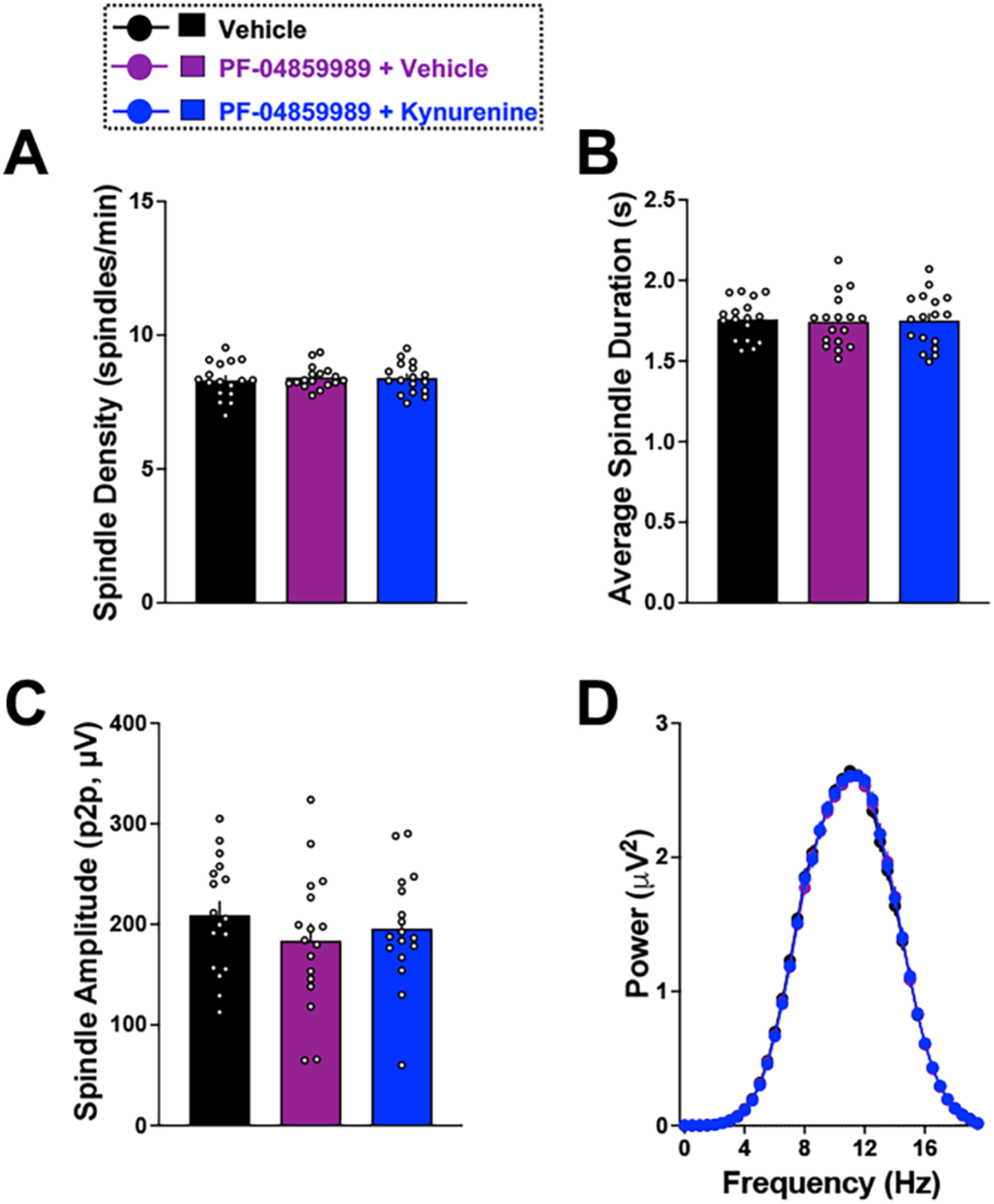
Pretreatment with PF-04859989 prior to kynurenine challenge restores NREM spindle dynamics. Adult rats were treated with vehicle, PF-04859989 (30 mg/kg), or PF-04859989 + kynurenine (100 mg/kg) at the start of the light phase. Sleep spindles were evaluated from ZT0-4. Data are mean ± SEM, analyzed by RM ANOVA. **(A)** NREMS spindle density (spindles/min NREMS). *F*_2,32_ = 0.2635, P=0.7700. **(B)** Average spindle duration. *F*_2,32_ = 0.1742, P=0.8409. **(C)** Spindle peak to peak (p2p) amplitude. *F*_2,32_ = 1.953, P=0.1583. **(D)** NREMS spindle power. Frequency: *F*_39,663_ = 840.3, P<0.0001. N = 18 per group (9 males, 9 females).

### PF-04859989 prevents kynurenine-induced changes in wake behavior

We found no significant differences in wake duration between Sleep Study #2 treatment conditions when evaluating data in 1-hr bins (**Supplemental Figure 6A**) or 6-hr bins (**Supplemental Figure 6B**). Wake bout number (**Supplemental Figure 6C**) and average duration of each wake bout (**Supplemental Figure 6D**) were also not altered by PF-04859989 treatment. As in Sleep Study #1 home cage activity data, we similarly evaluated activity separately by sex in Sleep Study #2. Home cage activity was significantly impacted by an interaction between ZT and treatment in males, with increased activity at ZT 1 (PF-04859989 + vehicle: P<0.001, PF-04859989 + kynurenine: P<0.001), and reduced activity at ZT 4 (PF-04859989 + kynurenine: P<0.05) and ZT 24 (PF-04859989 + vehicle: P<0.001, PF-04859989 + kynurenine: P<0.01) (**Supplemental Figure 6E**). PF-04859989 treatment with or without kynurenine did not change home cage activity in female subjects.

### Vigilance state transitions are not impacted by PF-04859989 treatment

PF-04859989 successfully rescued kynurenine-induced changes in sleep transitions (**Supplemental Table 4**), as in Sleep Study #2 no significant differences in transitions were noted between treatment conditions. Bout length distribution analysis in the light phase showed no impact of PF-04859989 treatment (**Supplemental Figure 7**).

### PF-04859989 treatment attenuates sleep disruptions induced by kynurenine challenge

Vigilance state durations, sleep spindle density, and NREMS slow wave spectral power were normalized to a percent change from vehicle treatment to compare results from Sleep Study #1 with Sleep Study #2. Kynurenine challenge resulted in a 11% decrease in REMS duration and this impact was significantly attenuated with PF-04859989 + kynurenine treatment (P<0.0001)(**Figure 6A**). PF-04859989 given prior to kynurenine challenge significantly attenuated the reduction in slow wave (0.5 – 2 Hz) spectral power during NREMS with kynurenine challenge alone (**Figure 6B)**. Lastly, spindle density was reduced by 12% with kynurenine challenge and PF-04859989 + kynurenine treatment substantially rescued this phenotype (P<0.01)(**Figure 6C)**. Taken together, we demonstrate the ability of KAT II inhibitor PF-04859989 to reverse kynurenine-induced sleep spindle and REMS abnormalities (**Figure 6**).

**Figure 6.**
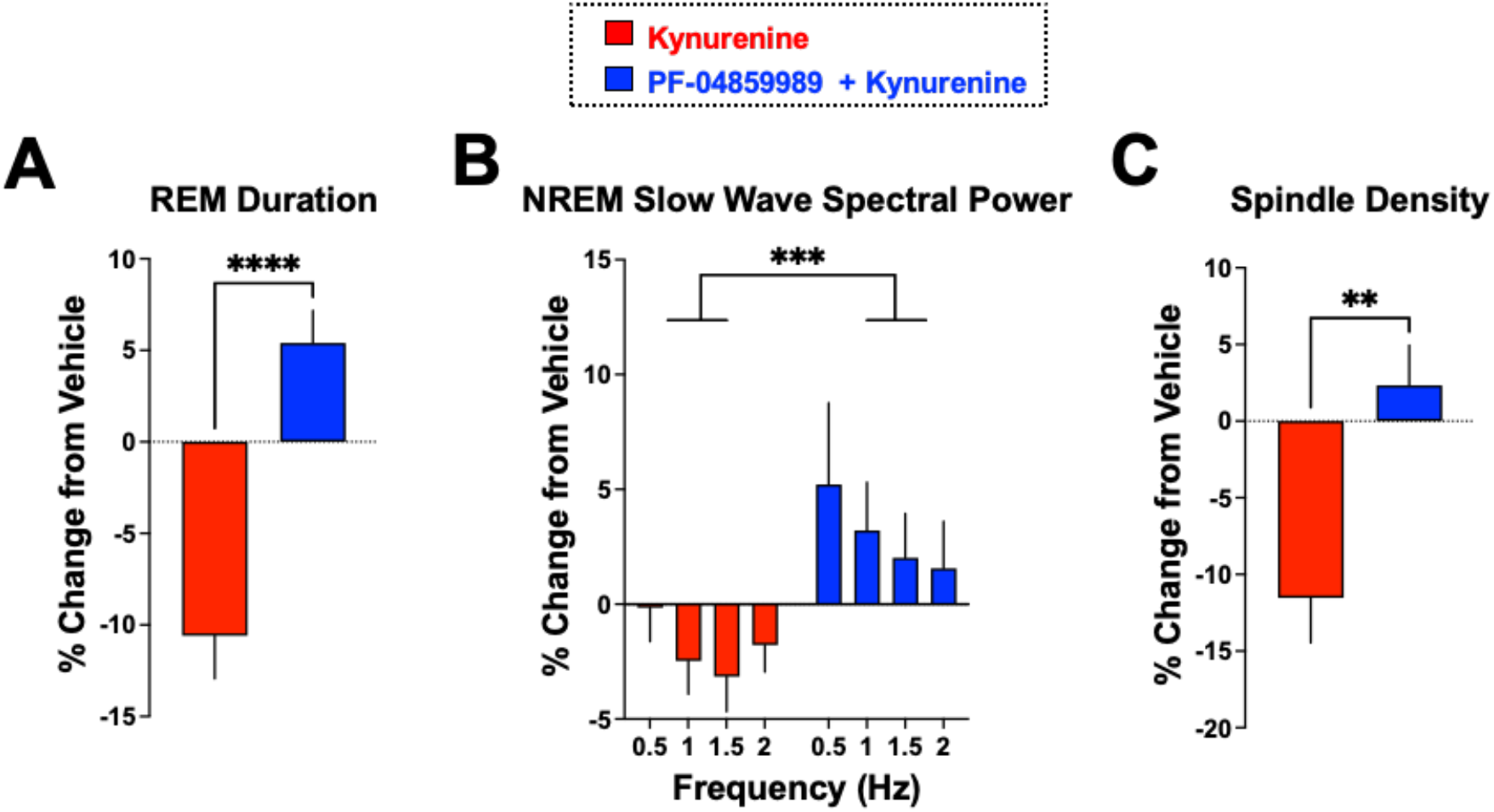
Pretreatment with PF-04859989 attenuates consequences of kynurenine challenge on REM and NREM sleep parameters. In Sleep Study #1, adult rats were treated with vehicle or kynurenine (100 mg/kg) at Zeitgeber time (ZT) 0. In Sleep Study #2, adult rats were treated with vehicle (ZT 23.5 and ZT 0), PF-04859989 (30 mg/kg; ZT 23.5) + vehicle (ZT 0) or PF-04859989 (30 mg/kg; ZT 23.5) + kynurenine (100 mg/kg; ZT 0). To compare results from the two sleep studies, data were evaluated as percent change from vehicle. **(A)** REMS duration (ZT 0-12); Student’s T test, t_36_ = 5.344, ****P<0.0001 **(B)** NREMS slow wave spectral power (ZT 0-12), two-way ANOVA, *F*_1,144_ = 12.31, ***P<0.001 **(C)** spindle density (ZT 0-4); Student’s T test, t_36_ = 3.495, **P<0.01. All data are mean ± SEM. Kynurenine: N = 20; PF-04859989 + Kynurenine: N = 18.

## DISCUSSION

Altered tryptophan metabolism, including KP dysregulation, impacts sleep and arousal in normal and pathological conditions (Milosavljevic et al., 2023; Mukherjee et al., 2018; Pocivavsek et al., 2017; Rentschler et al., 2021; Yamashita, 2020). Using telemetric recordings for the acquisition of EEG, EMG, and activity data in freely moving animals, we obtained ethologically relevant polysomnography recordings to evaluate sleep-wake behavior with acute elevation of KYNA (kynurenine challenge) or inhibition of newly produced KYNA in the brain (PF-04859989 + kynurenine). Building upon previous work, which evaluated changes in sleep with a dose-response of kynurenine challenge in male rats (Pocivavsek et al., 2017), we presently expanded our understanding of the impact on sleep-wake behaviors by inhibiting KAT II, thereby reducing de novo KYNA formation, with a brain-penetrable and irreversible compound, PF-04859989 in male and female subjects.

Acute kynurenine challenge notably reduced the total duration of REMS and the architecture of sleep, denoted as reduced REMS bouts, time bin distribution of sleep bouts, and transitions from NREMS to REMS and vice versa. Changes in sleep-wake architecture induced by kynurenine challenge were prevented with prophylactic PF-04859989 treatment, and we observe that the reduced KYNA levels across various brain regions contributed to the attenuated sleep behavior phenotypes. Namely, PF-04859989 effectively reduced KYNA levels in the hypothalamus, basal forebrain, hippocampus, cortex, and frontal cortex, suggesting that the majority of KYNA synthesis in these regions is KAT II-dependent. While we saw slight trends towards reduced KYNA levels in the cerebellum and brainstem with PF-04859989 treatment prior to kynurenine challenge, the effects were not significant thereby endorsing evidence of alternative means to regulate KYNA formation within these brain regions (Blanco Ayala et al., 2021). At the transcriptome level, KAT II mRNA encoded by *Aadat* are found prominently in cerebral regions such as the hippocampus, and much less in cerebellum and brainstem (Song et al., 2018). In the case of the cerebellum, KYNA formation may be more dependent on non-enzymatic formation with a role for reactive oxygen species, and a peripheral KAT II inhibitor may be less effective (Blanco Ayala et al., 2021).

The basal forebrain, a region wherein KYNA was both elevated with kynurenine challenge and reduced with PF-04859989 pre-treatment, is richly implicated in modulation of sleep and arousal behavior (Lelkes, Abdurakhmanova, & Porkka-Heiskanen, 2018; Szymusiak, Alam, & McGinty, 2000). A large amount of acetylcholine is released from the basal forebrain during REM sleep (Jing et al., 2020), and we postulate that KYNA-mediated modulation of cholinergic neurotransmission may impede these dynamics (Zmarowski et al., 2009). Within the extracellular milieu, KYNA is present within low micromolar concentrations and optimally positioned to inhibit receptor targets including α7nACh receptors and NMDA receptors (Albuquerque & Schwarcz, 2013; Konradsson-Geuken et al., 2010). The stark decrease in REMS with acute KYNA elevation is presumably related to the inhibition of these targets. The fact that systemic application of PF-04859989 downregulated KYNA production and reversed the consequences of KYNA elevation on sleep behavior provides compelling evidence for the role of KYNA, rather than the alternate arm of the kynurenine pathway that produces the metabolites 3-hydroxykynureine, 3-hydroxyanthranilic acid, and quinolinic acid (Schwarcz et al., 2012), in modulating changes in sleep physiology. Ongoing work with mouse models genetically modified to eliminate key enzymes of the kynurenine pathway will provide further support for the implicated role of KYNA in sleep behavior (Erhardt et al., 2017).

Key sleep disruptions bring further attention to the concept that KYNA serves as a molecular intersection between sleep and cognitive function (Pocivavsek et al., 2017; Pocivavsek et al., 2011). Reduced NREMS duration, NREMS delta spectral power and disrupted transitions between NREMS and REMS with elevated KYNA may influence long-term memory consolidation (Tartar et al., 2006). Temporally, NREMS changes occurred during the early light phase and REMS duration and architecture, which impacts synaptic plasticity and memory (Boyce, Glasgow, Williams, & Adamantidis, 2016; Tartar et al., 2006), were subsequently impacted in the late light phase. Sleep spindles, oscillations produced by the thalamic reticular nucleus, support memory consolidation (Fernandez & Luthi, 2020) and are speculated to predict onset of REMS (Bandarabadi et al., 2020). Reduced spindle density is a hallmark endophenotype of sleep dysfunction in patients with psychotic disorders and considered a predictor for cognitive dysfunction (D’Agostino et al., 2018; Ferrarelli & Tononi, 2017; Manoach et al., 2014). Preclinical models relevant to the study of neuropsychiatric illnesses have noted reduced sleep spindle density in rodents (Aguilar, Strecker, Basheer, & McNally, 2020; Thankachan et al., 2019). Importantly, we show for the first time that a reduction of brain-derived KYNA, achieved with prophylactic PF-04859989 treatment, effectively corrected for kynurenine-induced spindle density deficiencies and REMS disturbances.

It is noteworthy that PF-04859989 precluded the reduced NREMS delta power induced by kynurenine treatment alone, suggesting that reducing KYNA accumulation improves the quality of slow wave sleep (Lundahl, Deacon, Maurice, & Staner, 2012). NREMS quality is directly correlated with improved neuropsychological functioning and remains a physiological point of therapeutic interest for cognitive impairment associated with insomnia, neuropsychiatric disorders, and aging (Wilckens, Ferrarelli, Walker, & Buysse, 2018). It is plausible that KYNA modulates NREMS by influencing γ-aminobutyric acid (GABA) neurotransmission, as evidenced in microdialysis studies wherein elevated KYNA associates with reduced local extracellular GABA levels in rats (Beggiato et al., 2014; Wright et al., 2021). GABA receptor agonists are commonly prescribed for insomnia and NREM sleep disorders (Lankford et al., 2008; Walsh et al., 2011), yet these drugs, including benzodiazepines or zolpidem, carry a risk of drug dependence and side effects including prolonged sleepiness, delirium, aggressivity, and complex behaviors such as sleepwalking (Earl & Van Tyle, 2020). Our findings place new attention on KAT II inhibitors to modulate sleep quality, improve daytime functioning and arousal. Complementary experiments wherein we consider the use of KAT II inhibitors at different times of day and apply chronobiology to therapeutic approaches may have significant translational value to the treatment of sleep disturbances (Milosavljevic et al., 2023). Lastly, despite a number of disruptions in sleep parameters, we did not find major changes in arousal phenotypes with increased KYNA. However, enhanced home cage activity with PF-04859989 immediately after treatment warrants further investigation into its short term efficacy in attentional tasks, as KAT II inhibitors have been shown to acutely enhance cognition (Klausing et al., 2020; Pocivavsek et al., 2011). Lastly, unanswered questions about microarousals and local sleep events could conceivably be addressed with technical advancements, including in automatic approaches to sleep scoring in future studies (Rayan et al., 2022; Vyazovskiy et al., 2011).

## CONCLUSIONS

In summary, our findings indicate an important association between KYNA, the KAT II enzyme, and sleep-wake behavior. Our data suggest that pharmacological agents that inhibit KAT II activity in the brain may attenuate sleep disturbances. KYNA is considered a modulator of cognitive and sleep mechanisms, thereby KYNA may causally disrupt the relationship between sleep and cognition in pathological conditions. Sleep dysfunction and cognitive impairments are associated with a variety of neurocognitive disorders (Brzecka et al., 2018; Manoach et al., 2014), and similarly elevated KYNA levels are found in the brains of patients with these conditions (Linderholm et al., 2012; Miller et al., 2006; Sathyasaikumar et al., 2011). As such, regulating KYNA levels with KAT II inhibition should be carefully considered in the development of novel therapeutic strategies to treat sleep and cognitive disturbances (Pocivavsek & Erhardt, 2023), and these strategies may provide value to improving the quality of life in patients afflicted with neurocognitive disorders and associated sleep disruptions.

## Acknowledgements

The authors would like to thank Julie Fox and Nathan TJ Wagner for their technical contributions to this work. This research was supported by National Institutes of Health Grant Nos. R01 NS102209 (AP), P50 MH103222 (AP), and in part by National Science Foundation under Grant No. 185233 and the University of South Carolina Office of the Vice President for Research.

The authors would like to thank Julie Fox and Nathan TJ Wagner for their technical contributions to this work.

## SUPPLEMENTAL METHODS

### Vigilance state scoring

The image below demonstrates representative hypnograms indicating Wake, REMS, or NREMS transitions across the light phase, ZT 0-12. The red circle denotes wherein we provide an expanded views of the continuous recording of EEG, EMG, and activity visualized in NeuroScore software. Representative examples of transitions from REMS to NREMS, NREMS to REMS, and REMS to Wake are provided.

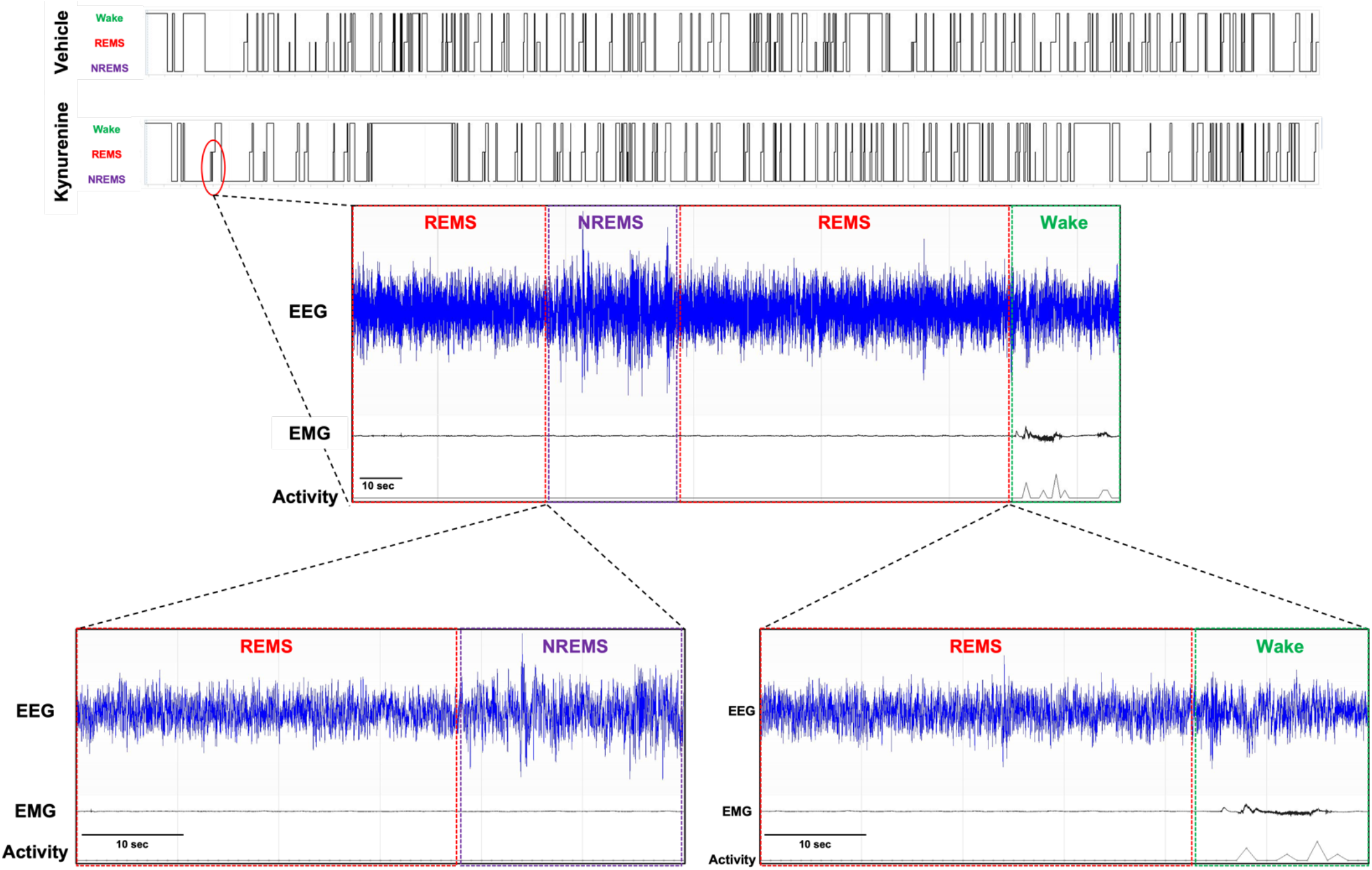

### Spindle Detection

We automatically detected sleep spindles using an approach similar to Uygun et al. (2019) proposed for spindle detection in mice. In short, EEG was first band-pass filtered (10 - 15 Hz; default parameters used by mne.io.Raw.filter) and resampled to 50 Hz to reduce unnecessary computation. We then computed the cube root-mean-square amplitude using a centered 0.75 s window sliding one sample at a time. This procedure gave us a spindle index signal x(n). For thresholding, we computed a baseline b(n) signal using the same approach but with a 600 s window. Using this slowly evolving rather than a constant baseline limits the effect of large artifacts to a 10-minute window and adjusts for any slow drift (e.g., slow changes in channel impedance). A low (1.2) and a high (3.5) threshold λ were used to define two sets of windows where x(n) > λ*b(n). The high threshold was used to detect spindles, whereas the low threshold was used to define the beginning and ending of the spindles. Only spindles detected with a duration between 0.5 s and 10 s were considered.

To validate the performance of this extractor, we computed the accuracy, recall, precision, and F1 metrics compared to manual scoring using a *signal-sample-based* approach (O’Reilly & Nielsen, 2015)(i.e., considering each sample from the EEG signal resampled at 50 Hz as a single false/true positive/negative) and reported these performances below. These performances are in the upper range of what can be expected for automated detectors where neither the expert (through training using information obtained from the automated detector) or the automated detector (i.e., through machine learning) were trained to match the behavior of one-another (Warby et al., 2014).

Briefly, manual validation in NeuroScore applied a rectified 10 - 15 Hz finite impulse response band pass filter to the EEG signal and computed the cubed root-mean square amplitude of the band pass filtered EEG signal within a 750 ms window to create a signal envelope for the accurate detection of sleep spindles. To ensure scientific rigor, all artifactual spikes with a duration of less than 0.5 s were excluded from spindle analyses criteria.

### Detection performance of the automated detection compared to expert scoring. Sample size (N) is for the number of 4 h (ZT0-ZT4) recording segments in the different experimental conditions

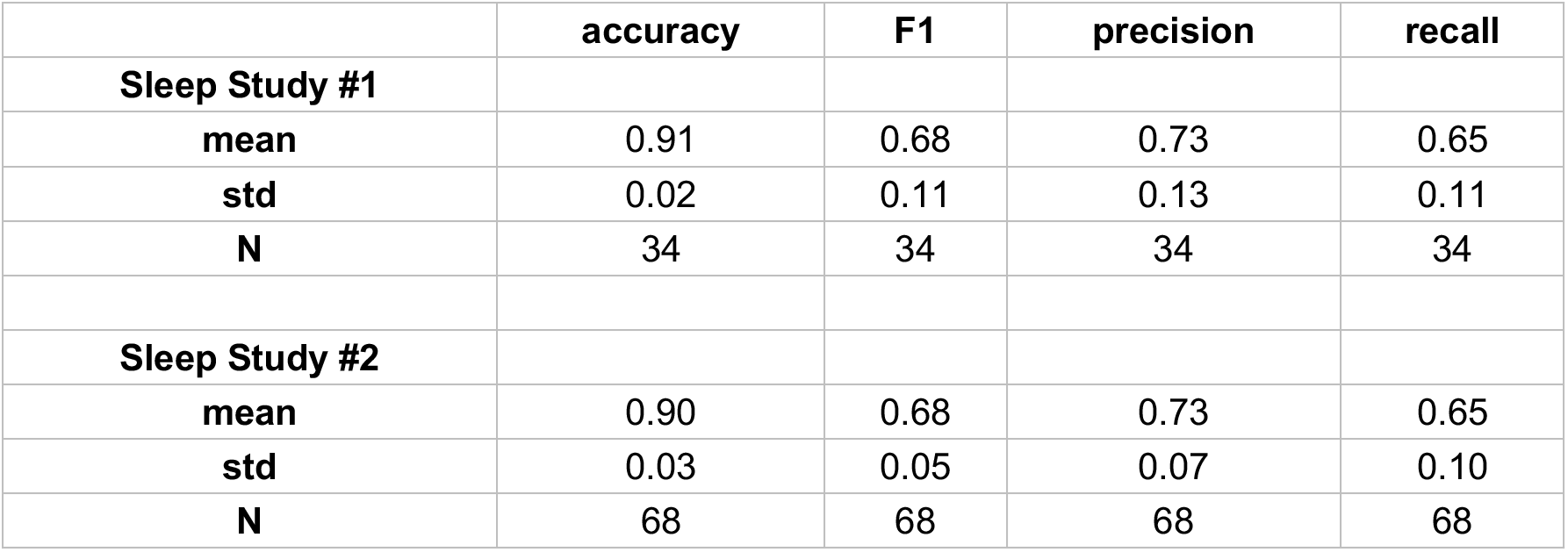

Spindle peak-to-peak amplitudes were computed identically for manual and automated detection by subtracting the maximum value from the minimum value of the 10 – 15 Hz band-pass filtered signals in the spindle windows. Densities were computed per 12-minute bins, dividing the number of spindles per the number of minutes of NREM sleep during these contiguous bins. The code for these analyses is available at https://github.com/lina-usc/paper-jsleepres-2023.

---

**Supplemental Figure 1.**
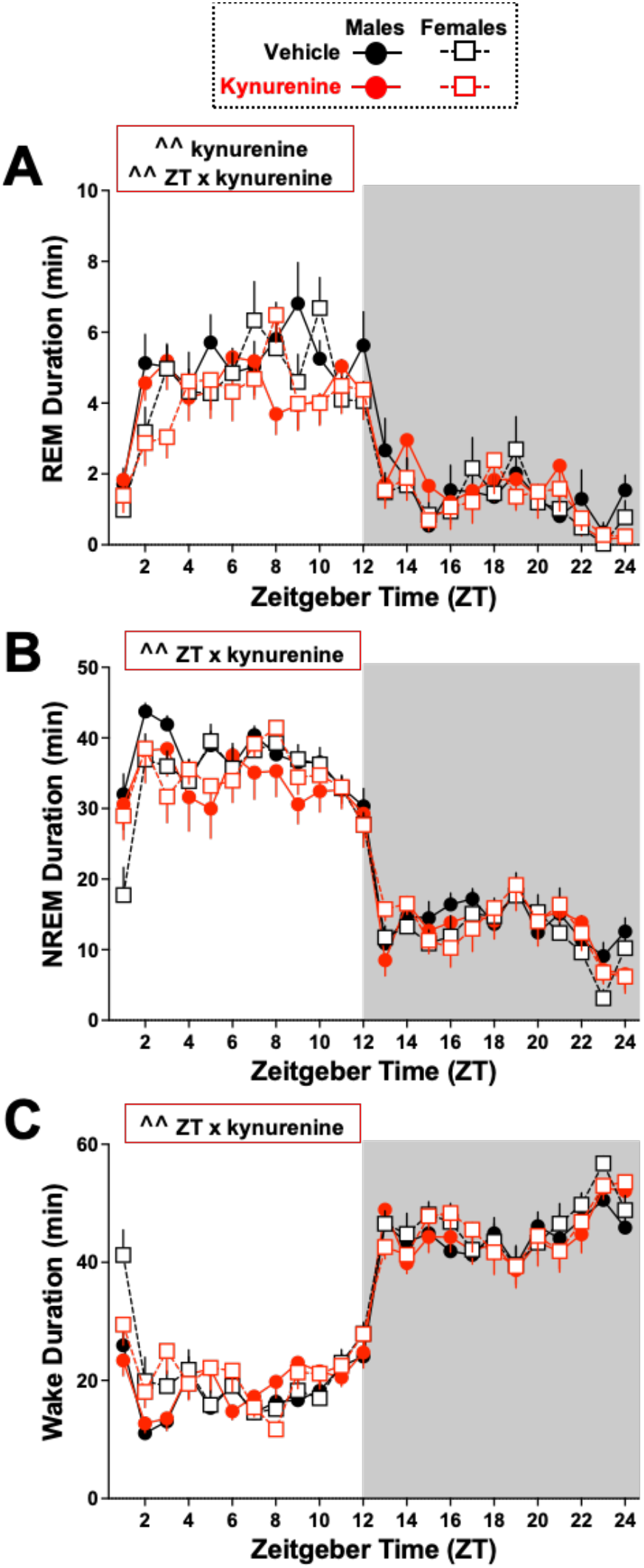
Impacts of kynurenine challenge on vigilance state duration. Adult rats were treated with vehicle or kynurenine (100 mg/kg) at Zeitgeber time (ZT) 0. Data are mean ± SEM, analyzed by three-way RM ANOVA, with significance shown in red boxes in graphs as ^^P<0.01. N= 8 males, 12 females. **(A)** 1-hr bins of REMS duration. ZT x Treatment: *F*_23,414_ = 1.618, P<0.01; Treatment: *F*_1,18_ = 15.44, P<0.01; ZT: *F*_23,414_ = 29.37, P<0.0001. Sex: *F*_1,18_ = 0.4279, P=0.5213. **(B)** 1-hr bins of NREMS duration. ZT x Treatment: *F*_23,414_ = 1.789, P<0.01 ZT: *F*_23,414_ = 66.12, P<0.0001. Sex: *F*_1,18_ = 0.3687, P=0.5513. **(C)** 1-hr bins of wake duration. ZT x Treatment: *F*_23,414_ = 1.807, P<0.01; ZT: *F*_23,414_ = 70.51, P<0.0001. Sex: *F*_1,18_ = 2.327, P=0.1445.

**Supplemental Figure 2.**
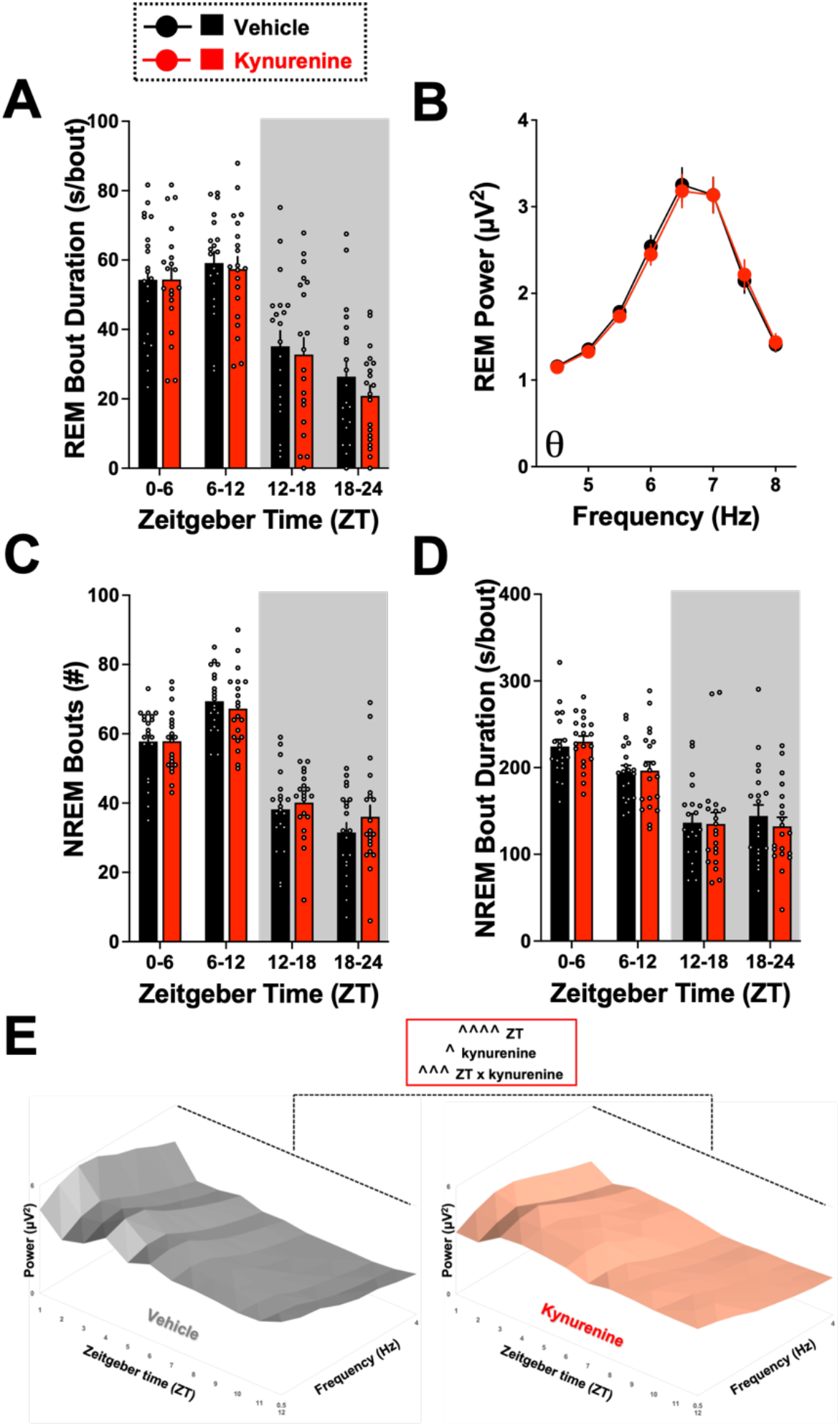
Impacts of kynurenine challenge on sleep architecture parameters. Adult rats were treated with vehicle or kynurenine (100 mg/kg) at Zeitgeber time (ZT) 0. Data are mean ± SEM, analyzed by two-way RM ANOVA. **(A)** 6-hr bins of average REMS bout duration. ZT: *F*_3,57_ = 48.65, P<0.0001. **(B)** REMS spectral power in the theta frequency range during the light phase. Frequency: *F*_7,133_ = 54.11, P<0.0001. **(C)** 6-hr bins of NREMS bout number. ZT: *F*_3,57_ = 75.97, P<0.0001. **(D)** 6-hr bins of average NREMS bout duration. ZT: *F*_3,57_ = 44.36, P<0.0001. **(E)** Delta spectra power (0.5 – 4 Hz) normalized to total spectra power visualized in 1-hr bins. ZT: *F*_11,187_ = 15.98, P<0.0001. Treatment: *F*_1,17_ = 4.615, P<0.05. ZT x treatment: *F*_11,187_ = 3.233, P<0.001. N = 20 per group (8 males, 12 females).

**Supplemental Figure 3.**
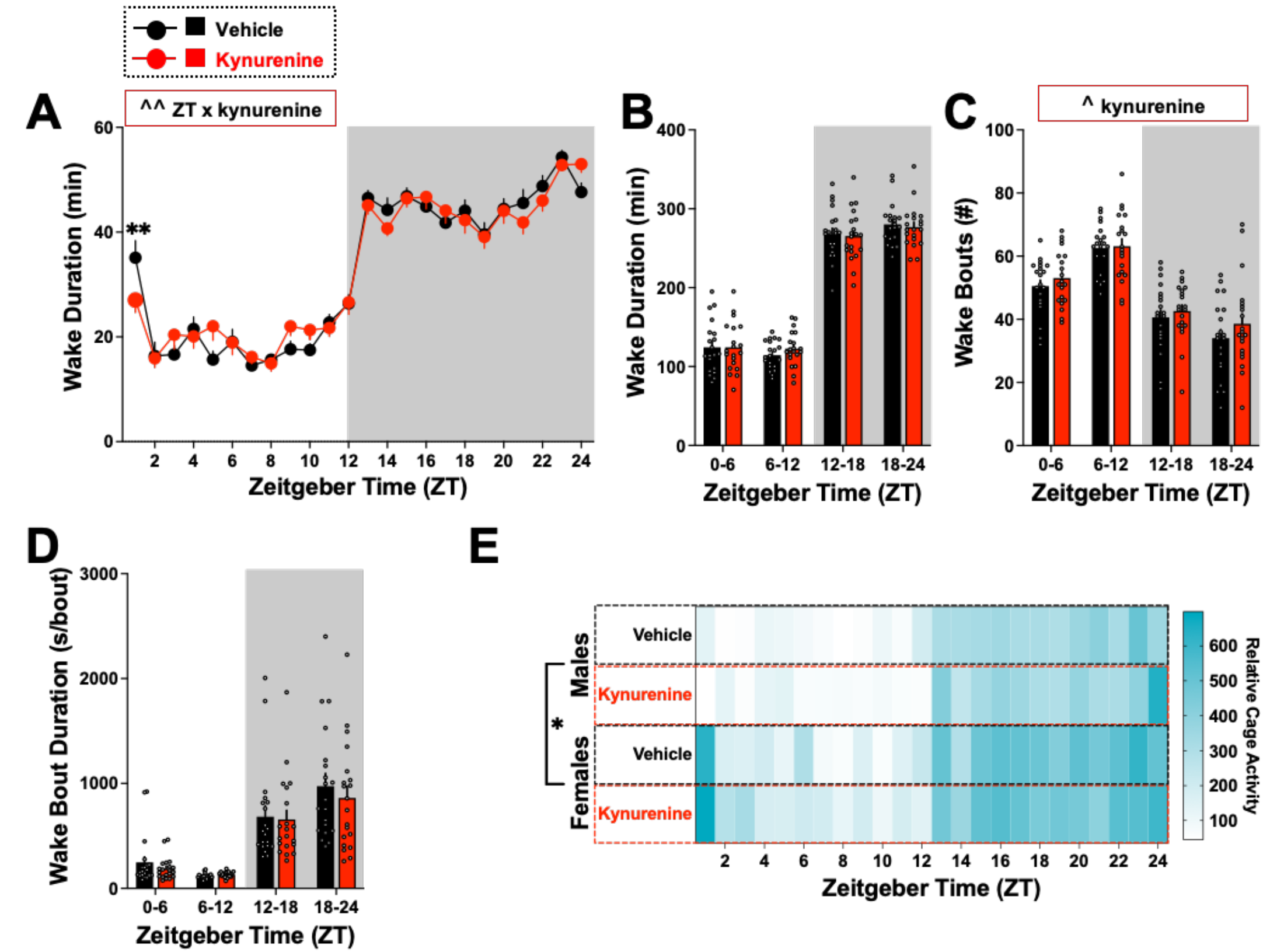
Impacts of kynurenine challenge on wake parameters. Adult rats were treated with vehicle or kynurenine (100 mg/kg) at Zeitgeber time (ZT) 0. Data are mean ± SEM, analyzed by two-way RM ANOVA (A-D), with significance shown in graphs as ^P<0.05, ^^P<0.01 and Bonferroni’s post hoc as **P<0.01 or three-way RM ANOVA (E), with significance shown in graph as *P<0.05. **(A)** 1-hr bins of wake duration. ZT x treatment: *F*_23,437_ = 2.078, P<0.01. ZT: *F*_23,437_ = 72.67, P<0.0001. **(B)** 6-hr bins of wake duration. ZT: *F*_3,57_ = 265.5, P<0.0001. **(C)** 6-hr bins of wake bout number. Treatment: *F*_1,19_ = 4.967, P<0.05. ZT: *F*_3,57_ = 44.76, P<0.0001. **(D)** 6-hr bins of average duration per wake bout. ZT: *F*_3,57_ = 34.07, P<0.0001. **(E)** Relative home cage activity. Sex: *F*_1,347_ = 5.118, *P<0.05. ZT: *F*_22,396_ = 25.40, P<0.0001. N = 20 per group (8 males, 12 females).

**Supplemental Figure 4.**
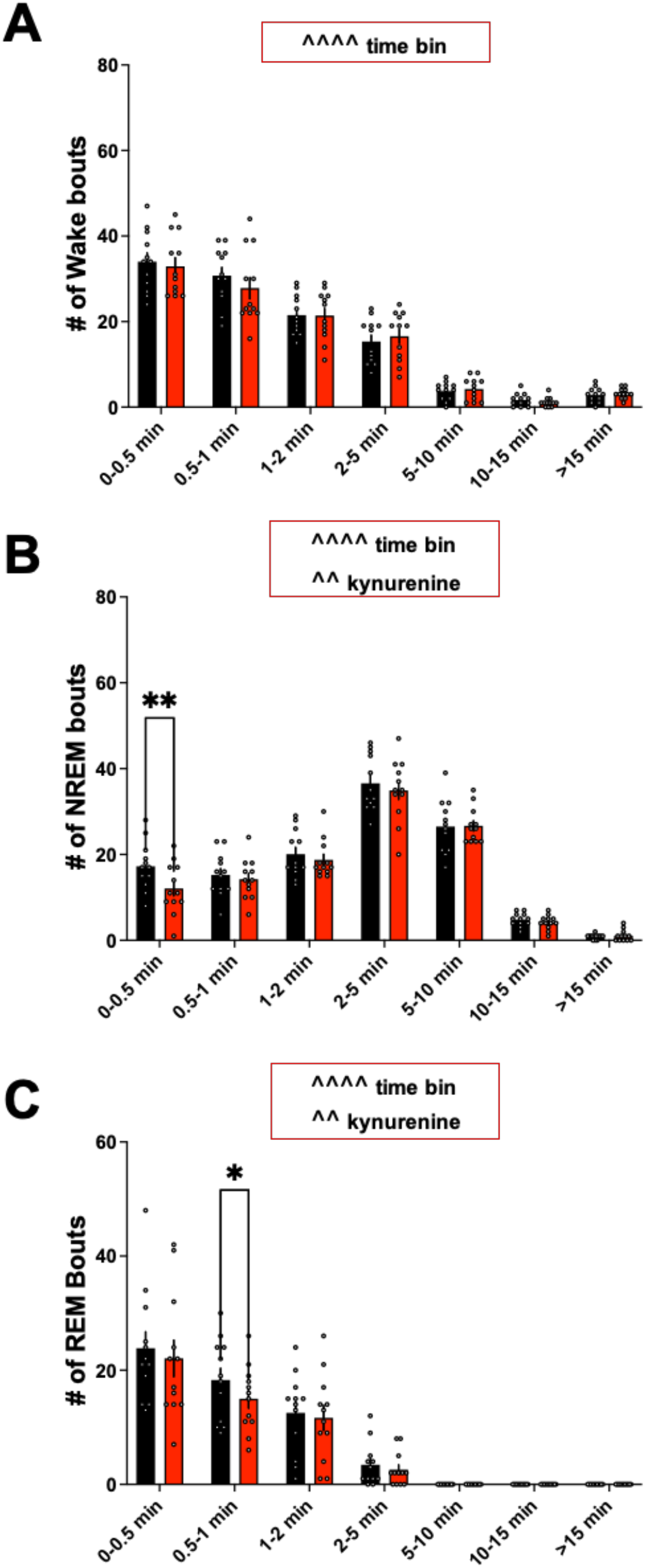
Kynurenine challenge significantly influenced NREM sleep and REM sleep bout length distribution. Adult rats were treated with vehicle or kynurenine (100 mg/kg) at Zeitgeber time (ZT) 0. Data are mean ± SEM, analyzed by two-way RM ANOVA (A-c), with significance shown in graphs as ^^P<0.01, ^^^^P<0.0001 and Bonferroni’s post hoc as *P<0.05, **P<0.01. **(A)** Time bins of wake bouts. Time bin: *F*_6,66_= 105.7, P<0.0001. **(B)** Time bins of NREM sleep bouts. Time bin: *F*_6,66_= 112.6, P<0.0001. Kynurenine: *F*_1,11_= 10.17, P<0.01. **(C)** Time bins of REM sleep bouts. Time bin: *F*_6,66_= 34.60, P<0.0001. Kynurenine: *F*_1,11_= 11.01, P<0.01. N = 12 per group (5 males, 7 females).

**Supplemental Figure 5.**
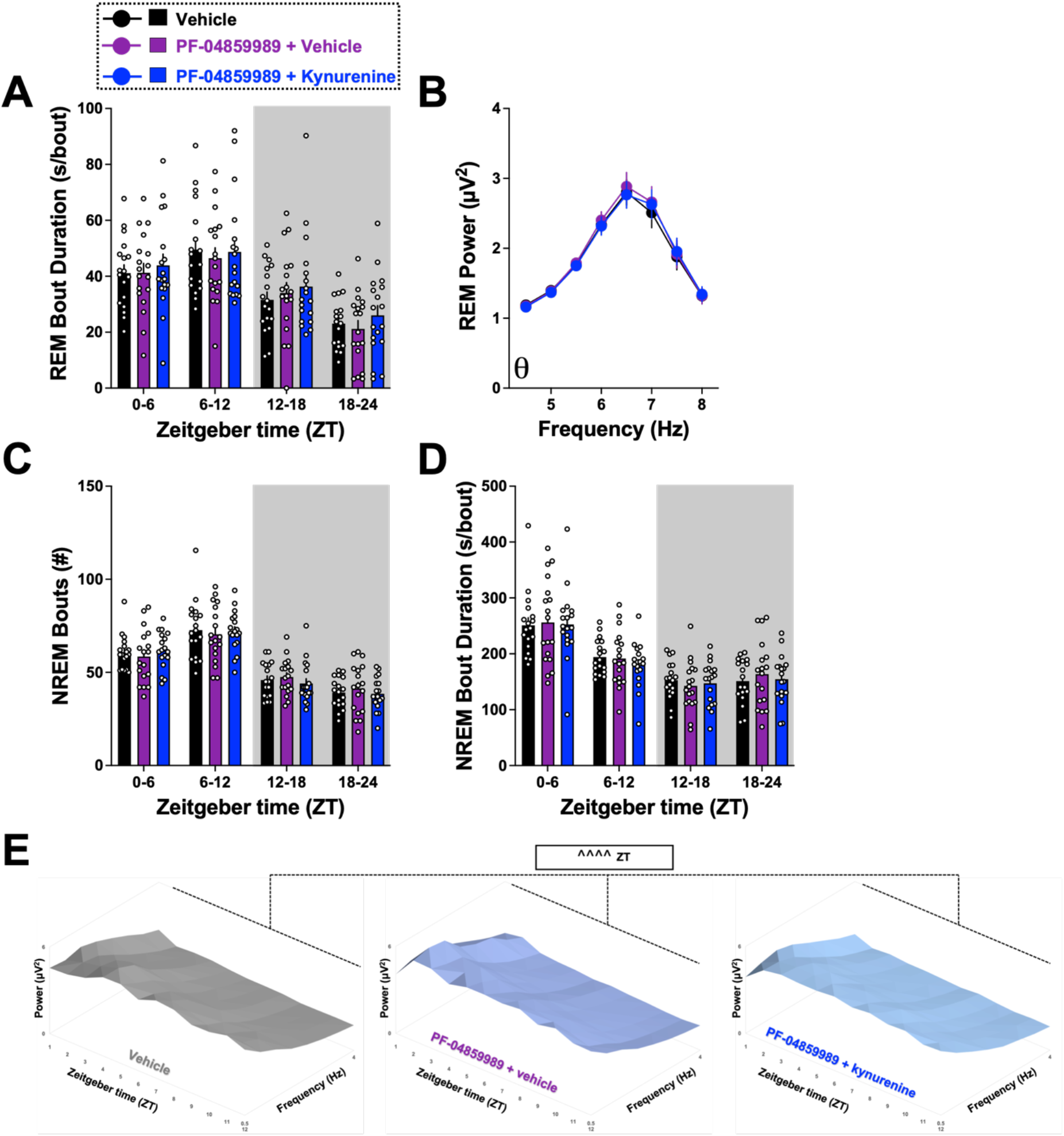
Impacts of PF-04859989 treatment prior to kynurenine challenge on sleep architecture parameters. Adult rats were treated with vehicle, PF-04859989 (30 mg/kg), or PF-04859989 + kynurenine (100 mg/kg) at the start of the light phase. Data are mean ± SEM, analyzed by RM ANOVA. **(A)** 6-hr bins of average REMS bout duration. ZT: *F*_3,51_ = 36.20, P<0.0001. **(B)** REMS spectral power in the theta frequency range during the light phase. Frequency: *F*_7,119_ = 29.01, P<0.0001. **(C)** 6-hr bins of NREMS bout number. ZT: *F*_3,51_ = 62.97, P<0.0001. **(D)** 6-hr bins of average NREMS bout duration. ZT: *F*_3,51_ = 68.22, P<0.0001. **(E)** Delta spectra power (0.5 – 4 Hz) normalized to total spectra power visualized in 1-hr bins. ZT: *F*_11,143_ = 10.80, P<0.0001. N = 18 per group (9 males, 9 females).

**Supplemental Figure 6.**
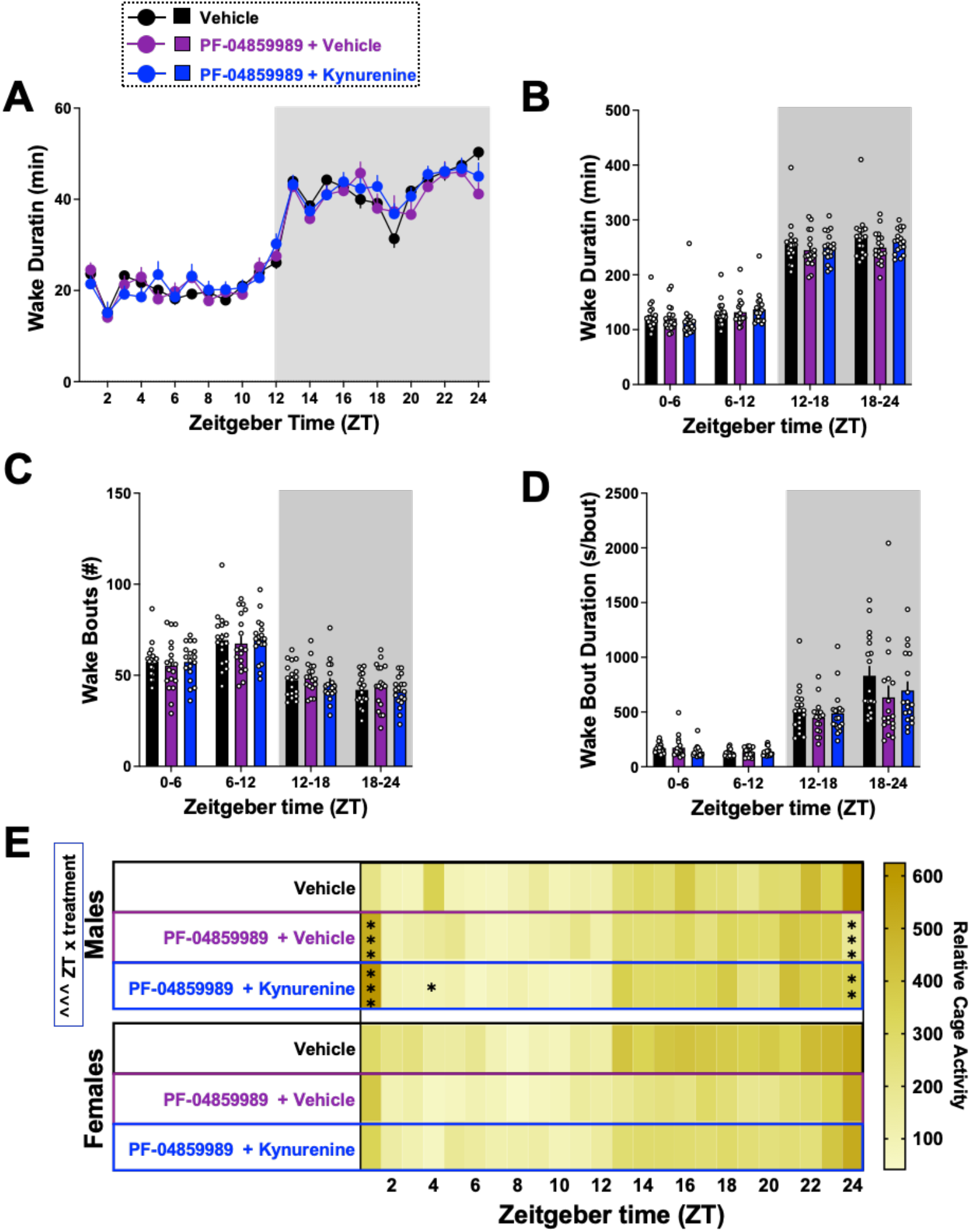
Impacts of pretreatment with PF-04859989 prior to kynurenine challenge on wake parameters. Adult rats were treated with vehicle, PF-04859989 (30 mg/kg), or PF-04859989 + kynurenine (100 mg/kg) at the start of the light phase. Data are mean ± SEM, analyzed by ANOVA, with significance shown in graphs as ^^^P<0.001, and Dunnett’s post hoc test *P<0.05, **P<0.01, ***P<0.001 vs vehicle. **(A)** 1-hr bins of wake duration. ZT: *F*_23,391_ = 87.96, P<0.0001. **(B)** 6-hr bins of wake duration. ZT: *F*_3,51_ = 283.1, P<0.0001. **(C)** 6-hr bins of wake bout number. ZT: *F*_3,51_ = 46.80, P<0.001. **(D)** 6-hr bins of average duration per wake bout. ZT: *F*_3,51_ = 51.67, P<0.0001. **(E)** Relative home cage activity, evaluated separated by sex. Males, ZT x treatment: *F*_46,322_ = 2.075, P<0.001; ZT: *F*_23,184_= 11.66, P<0.0001. Females, ZT: *F*_23,513_ = 13.83, P<0.0001. N = 18 per group (9 males, 9 females).

**Supplemental Figure 7.**
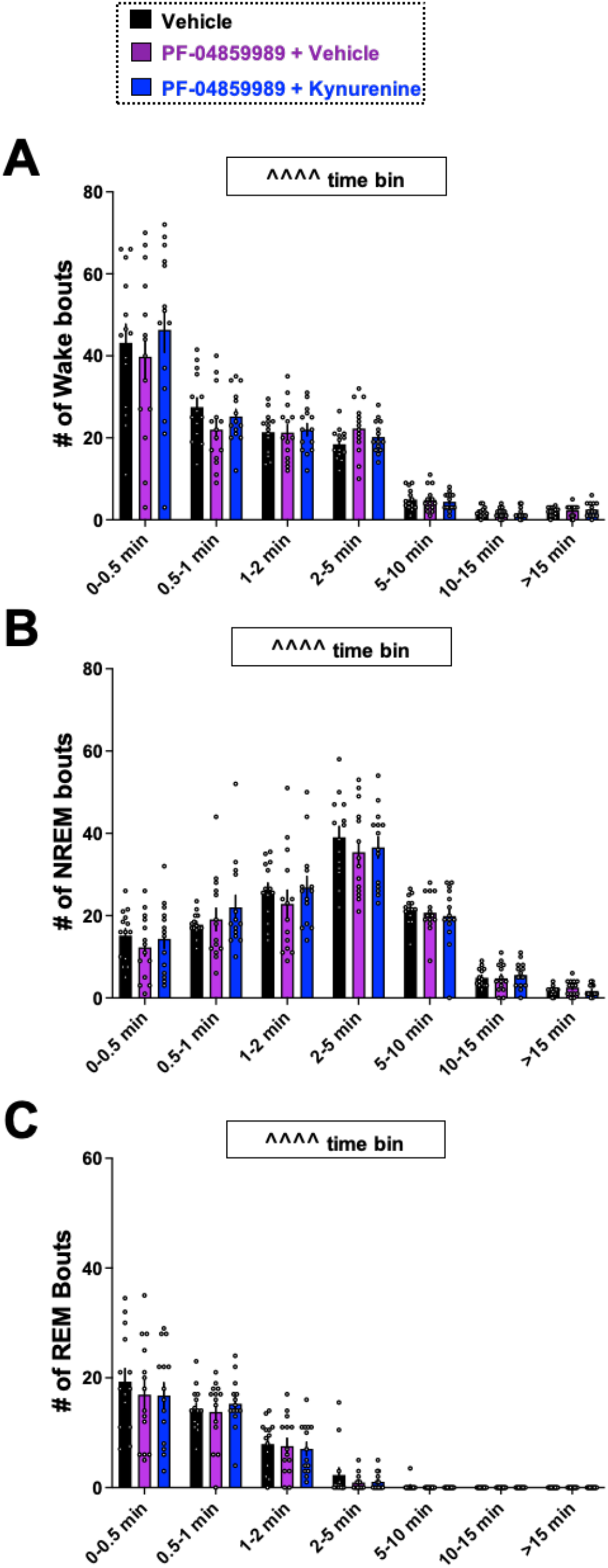
Impacts of pretreatment with PF-04859989 prior to kynurenine challenge on NREM sleep and REM sleep bout length distribution. Adult rats were treated with vehicle, PF-04859989 (30 mg/kg), or PF-04859989 + kynurenine (100 mg/kg) at the start of the light phase. Data are mean ± SEM, analyzed by ANOVA. **(A)** Time bins of wake bouts. Time bin: *F*_6,78_= 61.06, P<0.0001. **(B)** Time bins of NREM sleep bouts. Time bin: *F*_6,78_= 79.41, P<0.0001. **(C)** Time bins of REM sleep bouts. Time bin: *F*_6,78_= 55.26, P<0.0001. N = 14 per group (7 males, 7 females).

**Supplemental Table 1.**
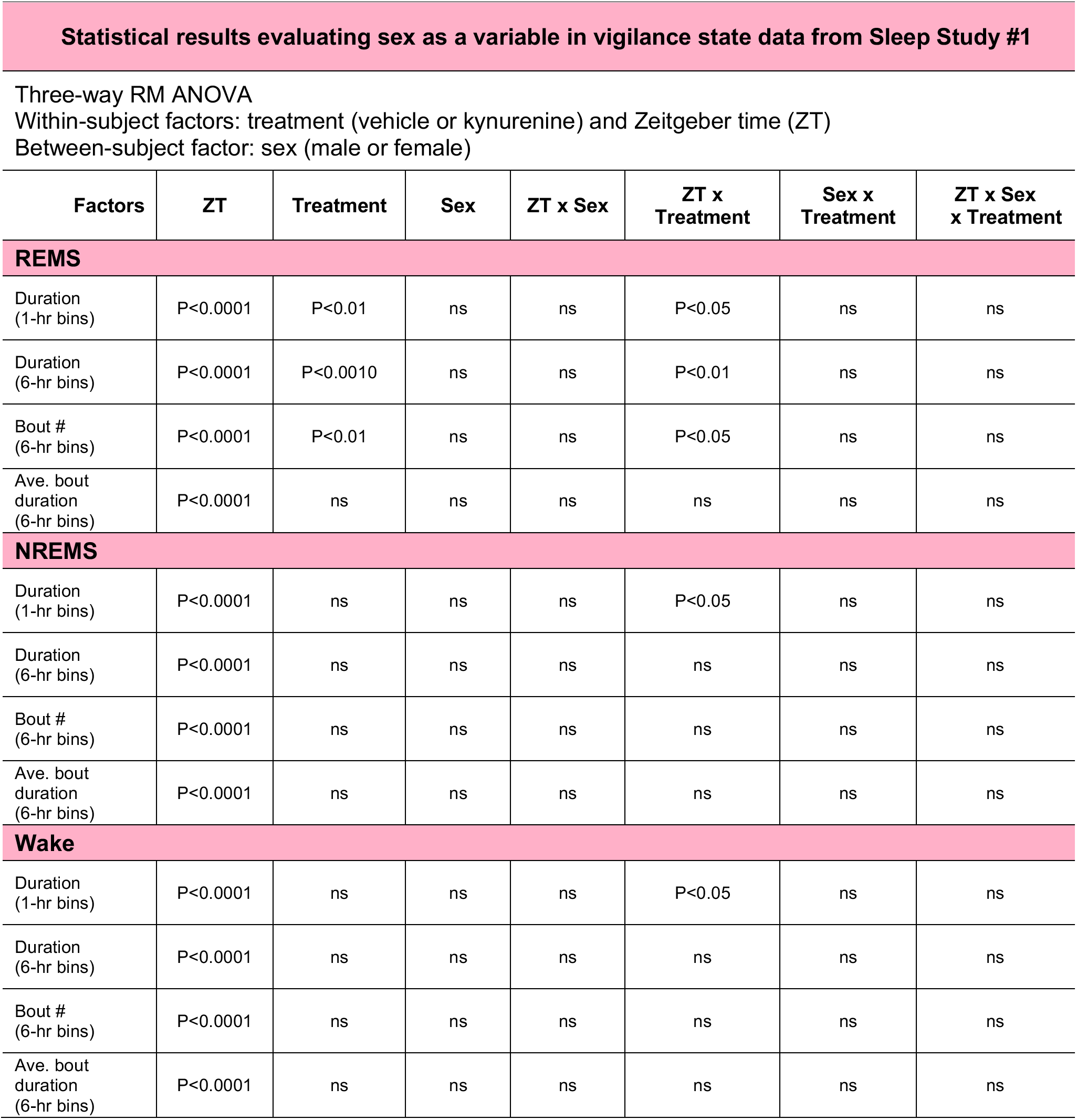
To evaluate sex as a biological variable, sleep architecture data (vigilance state durations, bout number, and average bout duration) were initially evaluated by 3-way repeated measures (RM) ANOVA with treatment and ZT as within-subject factors and sex as a between-subject factor. N = 20 per group (8 males, 12 females).

**Supplemental Table 2.**
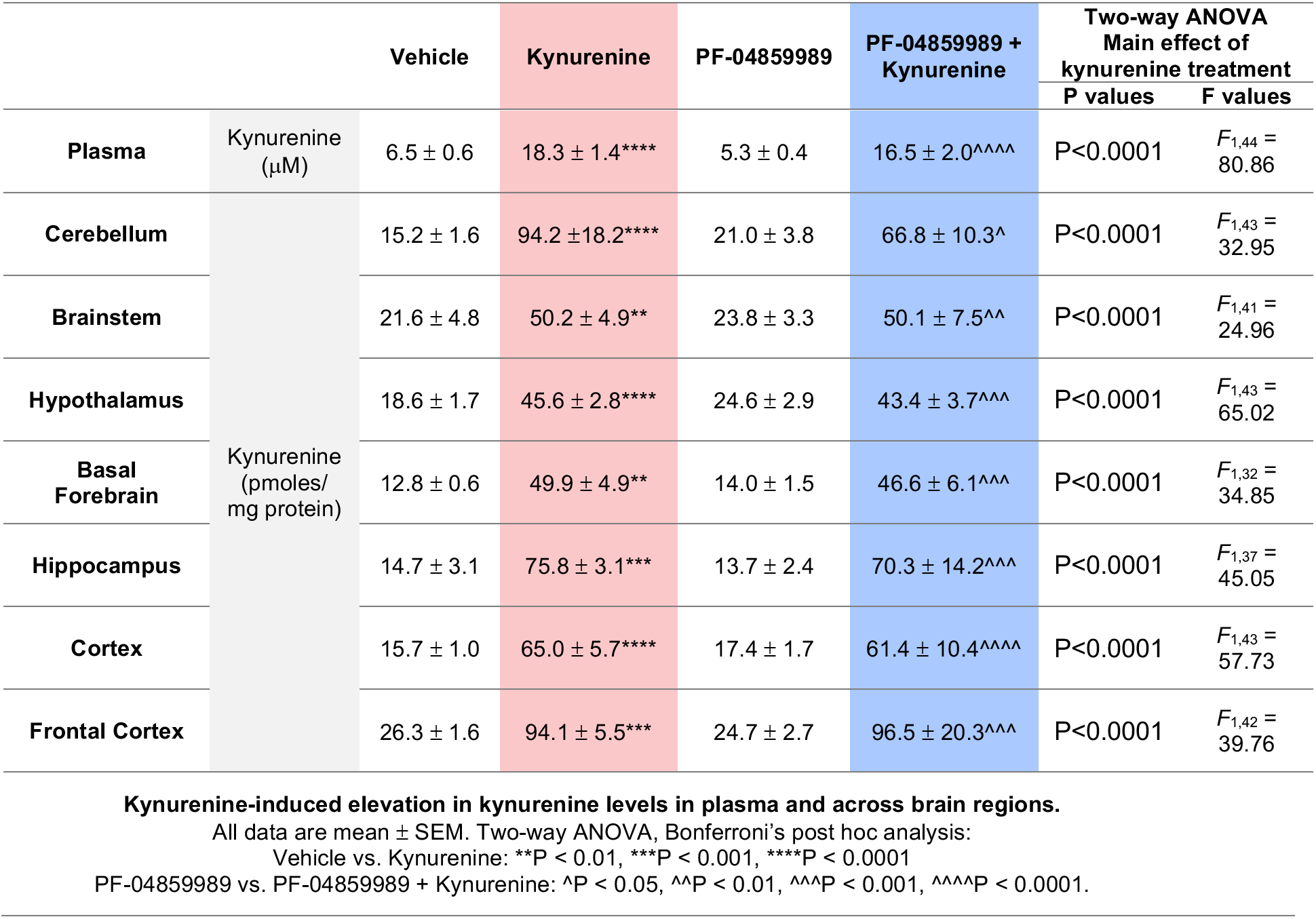
Kynurenine challenge significantly increases kynurenine levels in plasma and across brain regions. Adult rats were peripherally injected at Zeitgeber time (ZT) 0 with kynurenine (100 mg/kg) to induce de novo kynurenic acid (KYNA) formation. PF-04859989 (30 mg/kg), systemically active KAT II inhibitor, was given 30 minutes prior at ZT 23.5. Tissues were harvested at ZT 2. Kynurenine levels were evaluated. All data are mean ± SEM. N = 4 - 12 per group.

**Supplemental Table 3.**
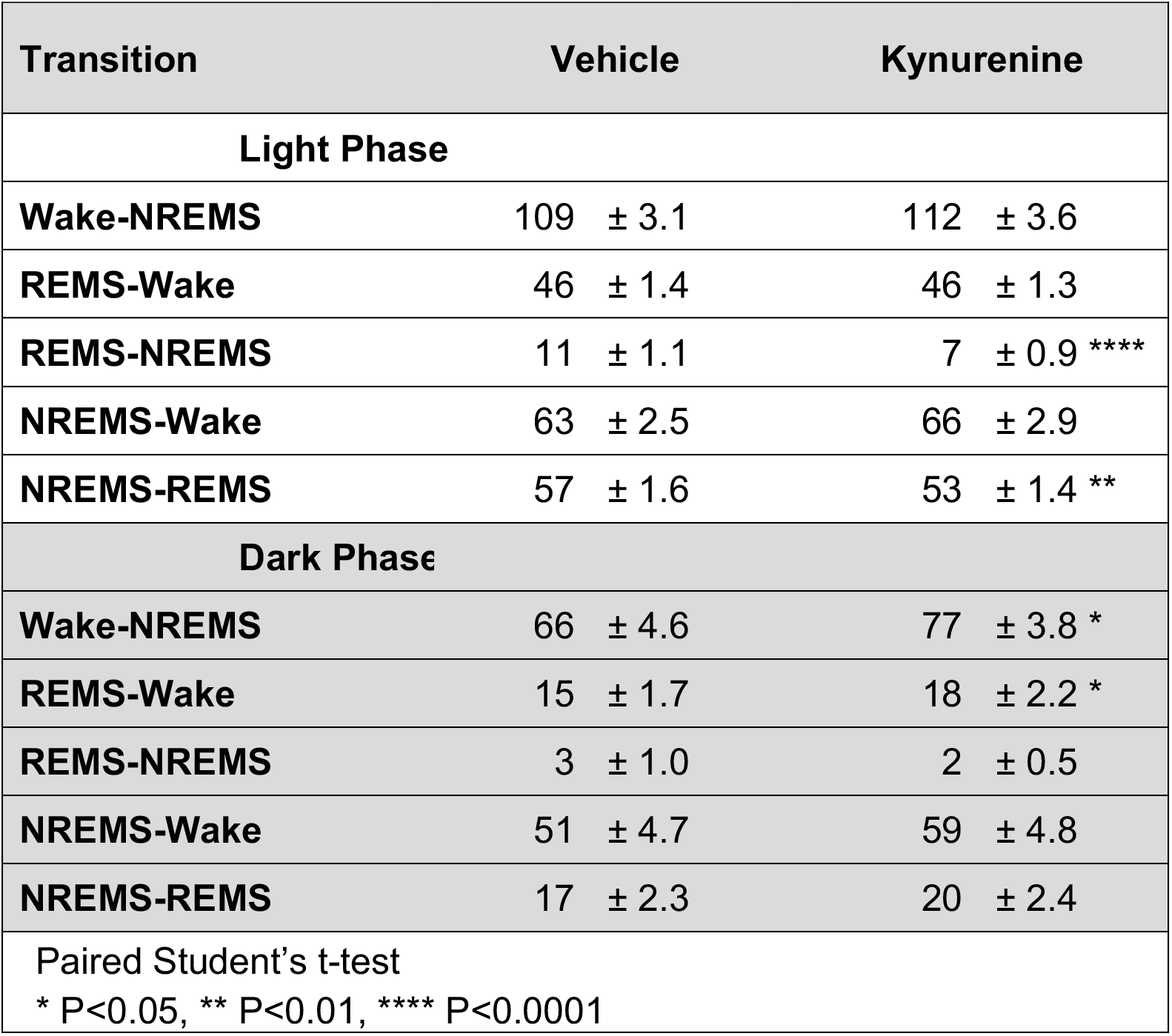
Acute kynurenine challenge disrupts female sleep transitions. Adult male and female rats were treated with vehicle or kynurenine (100 mg/kg) at Zeitgeber time (ZT) 0. All data are mean ± SEM, analyzed by paired Student’s t-test. N = 20 per group (8 males, 12 females).

**Supplemental Table 4.**
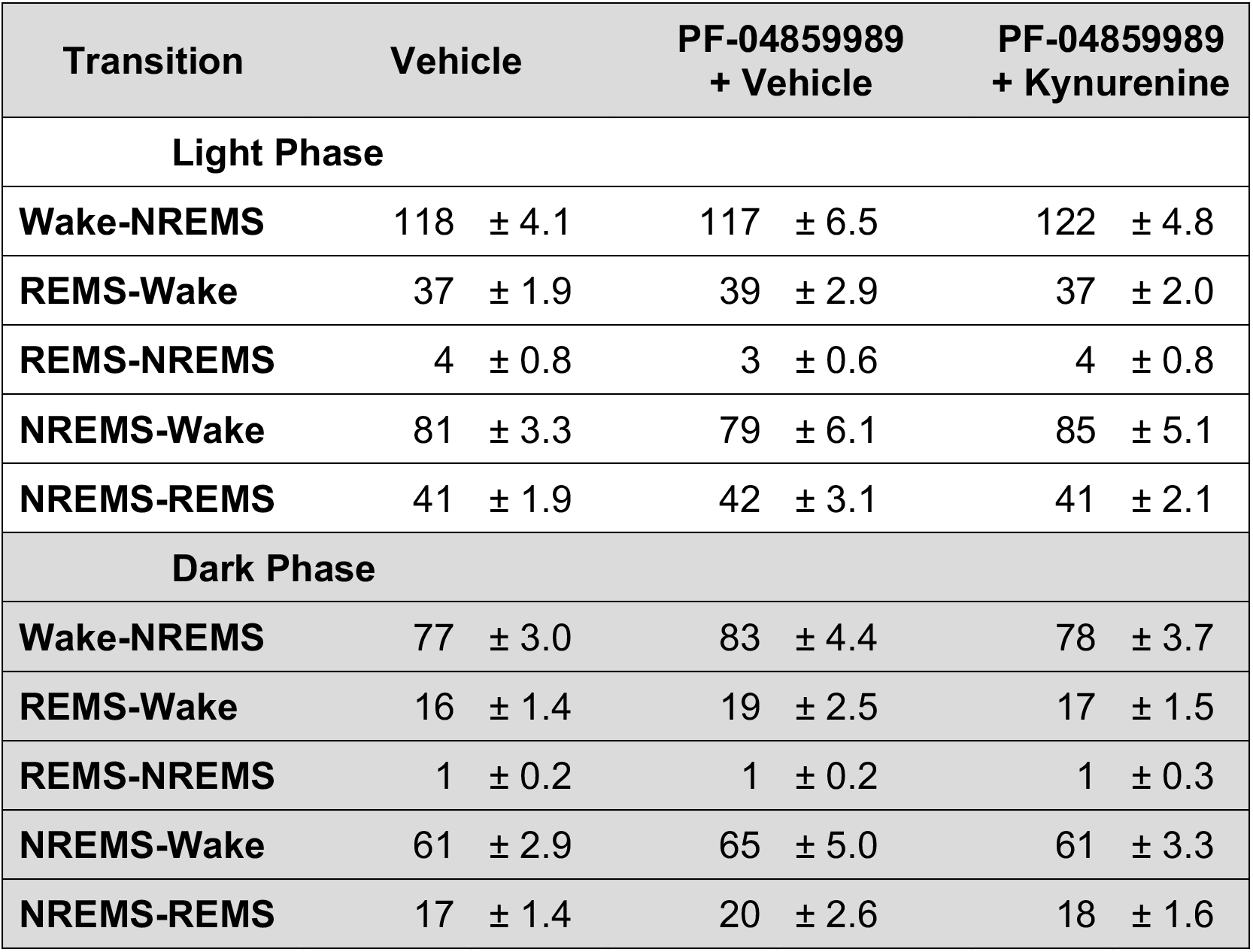
PF-04859989 treatment does not alter vigilance state transitions. Adult rats were treated with vehicle, PF-04859989 (30 mg/kg), or PF-04859989 + kynurenine (100 mg/kg) at the start of the light phase. All data are mean ± SEM, analyzed by ANOVA. N = 18 per group (9 males, 9 females).

