## Supplemental Materials for "Reducing brain kynurenic acid synthesis precludes kynurenine-induced sleep disturbances"

### SUPPLEMENTAL METHODS

**Vigilance state scoring.** The image below demonstrates representative hypnograms indicating Wake, REMS, or NREMS transitions across the light phase, ZT 0-12. The red circle denotes wherein we provide an expanded views of the continuous recording of EEG, EMG, and activity visualized in NeuroScore software. Representative examples of transitions from REMS to NREMS, NREMS to REMS, and REMS to Wake are provided.

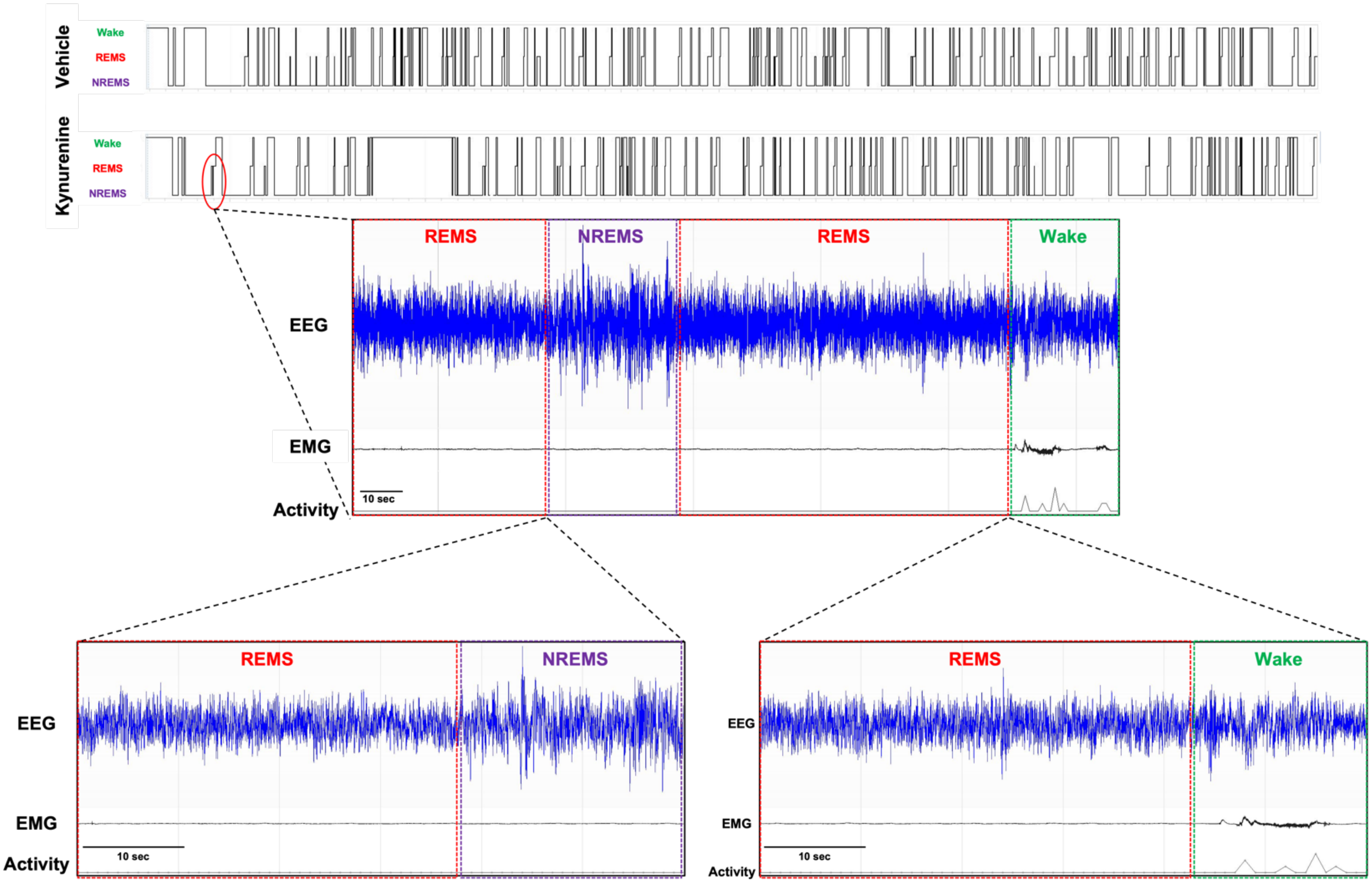

**Spindle Detection.** We automatically detected sleep spindles using an approach similar to Uygun et al. (2019) proposed for spindle detection in mice. In short, EEG was first band-pass filtered (10 - 15 Hz; default parameters used by mne.io.Raw.filter) and resampled to 50 Hz to reduce unnecessary computation. We then computed the cube root-mean-square amplitude using a centered 0.75 s window sliding one sample at a time. This procedure gave us a spindle index signal $x(n)$. For thresholding, we computed a baseline $b(n)$ signal using the same approach but with a 600 s window. Using this slowly evolving rather than a constant baseline limits the effect of large artifacts to a 10-minute window and adjusts for any slow drift (e.g., slow changes in channel impedance). A low (1.2) and a high (3.5) threshold $\lambda$ were used to define two sets of windows where $x(n) > \lambda * b(n)$. The high threshold was used to detect spindles, whereas the low threshold was used to define the beginning and ending of the spindles. Only spindles detected with a duration between 0.5 s and 10 s were considered.

To validate the performance of this extractor, we computed the accuracy, recall, precision, and F1 metrics compared to manual scoring using a *signal-sample-based* approach (O'Reilly & Nielsen, 2015)(i.e., considering each sample from the EEG signal resampled at 50 Hz as a single false/true positive/negative) and reported these performances below. These performances are in the upper range of what can be expected for automated detectors where neither the expert (through training using information obtained from the automated detector) or the automated detector (i.e., through machine learning) were trained to match the behavior of one-another (Warby et al., 2014).

Briefly, manual validation in NeuroScore applied a rectified 10 - 15 Hz finite impulse response band pass filter to the EEG signal and computed the cubed root-mean square amplitude of the band pass filtered EEG signal within a 750 ms window to create a signal envelope for the accurate detection of sleep spindles. To ensure scientific rigor, all artifactual spikes with a duration of less than 0.5 s were excluded from spindle analyses criteria.

**Detection performance of the automated detection compared to expert scoring. Sample size (N) is for the number of 4 h (ZT0-ZT4) recording segments in the different experimental conditions.**

| | **accuracy** | **F1** | **precision** | **recall** |
|---|---|---|---|---|
| **Sleep Study #1** | | | | |
| **mean** | 0.91 | 0.68 | 0.73 | 0.65 |
| **std** | 0.02 | 0.11 | 0.13 | 0.11 |
| **N** | 34 | 34 | 34 | 34 |
| **Sleep Study #2** | | | | |
| **mean** | 0.90 | 0.68 | 0.73 | 0.65 |
| **std** | 0.03 | 0.05 | 0.07 | 0.10 |
| **N** | 68 | 68 | 68 | 68 |

Spindle peak-to-peak amplitudes were computed identically for manual and automated detection by subtracting the maximum value from the minimum value of the 10 – 15 Hz band-pass filtered signals in the spindle windows. Densities were computed per 12-minute bins, dividing the number of spindles per the number of minutes of NREM sleep during these contiguous bins. The code for these analyses is available at https://github.com/lina-usc/paper-jsleepres-2023.

**Statistical results evaluating sex as a variable in vigilance state data from Sleep Study #1**

Three-way RM ANOVA
Within-subject factors: treatment (vehicle or kynurenine) and Zeitgeber time (ZT)
Between-subject factor: sex (male or female)

| Factors | ZT | Treatment | Sex | ZT x Sex | ZT x Treatment | Sex x Treatment | ZT x Sex x Treatment |
|---|---|---|---|---|---|---|---|
| **REMS** | | | | | | | |
| Duration (1-hr bins) | P<0.0001 | P<0.01 | ns | ns | P<0.05 | ns | ns |
| Duration (6-hr bins) | P<0.0001 | P<0.0010 | ns | ns | P<0.01 | ns | ns |
| Bout # (6-hr bins) | P<0.0001 | P<0.01 | ns | ns | P<0.05 | ns | ns |
| Ave. bout duration (6-hr bins) | P<0.0001 | ns | ns | ns | ns | ns | ns |
| **NREMS** | | | | | | | |
| Duration (1-hr bins) | P<0.0001 | ns | ns | ns | P<0.05 | ns | ns |
| Duration (6-hr bins) | P<0.0001 | ns | ns | ns | ns | ns | ns |
| Bout # (6-hr bins) | P<0.0001 | ns | ns | ns | ns | ns | ns |
| Ave. bout duration (6-hr bins) | P<0.0001 | ns | ns | ns | ns | ns | ns |
| **Wake** | | | | | | | |
| Duration (1-hr bins) | P<0.0001 | ns | ns | ns | P<0.05 | ns | ns |
| Duration (6-hr bins) | P<0.0001 | ns | ns | ns | ns | ns | ns |
| Bout # (6-hr bins) | P<0.0001 | ns | ns | ns | ns | ns | ns |
| Ave. bout duration (6-hr bins) | P<0.0001 | ns | ns | ns | ns | ns | ns |

**Supplemental Table 1**. To evaluate sex as a biological variable, sleep architecture data (vigilance state durations, bout number, and average bout duration) were initially evaluated by 3-way repeated measures (RM) ANOVA with treatment and ZT as within-subject factors and sex as a between-subject factor. N = 20 per group (8 males, 12 females).

| | | Vehicle | Kynurenine | PF-04859989 | PF-04859989 + Kynurenine | Two-way ANOVA Main effect of kynurenine treatment P values | F values |
|---|---|---|---|---|---|---|---|
| **Plasma** | Kynurenine (μM) | 6.5 ± 0.6 | 18.3 ± 1.4**** | 5.3 ± 0.4 | 16.5 ± 2.0^^^^ | P<0.0001 | $F_{1,44}$ = 80.86 |
| **Cerebellum** | Kynurenine (pmoles/ mg protein) | 15.2 ± 1.6 | 94.2 ±18.2**** | 21.0 ± 3.8 | 66.8 ± 10.3^ | P<0.0001 | $F_{1,43}$ = 32.95 |
| **Brainstem** | | 21.6 ± 4.8 | 50.2 ± 4.9** | 23.8 ± 3.3 | 50.1 ± 7.5^^ | P<0.0001 | $F_{1,41}$ = 24.96 |
| **Hypothalamus** | | 18.6 ± 1.7 | 45.6 ± 2.8**** | 24.6 ± 2.9 | 43.4 ± 3.7^^^ | P<0.0001 | $F_{1,43}$ = 65.02 |
| **Basal Forebrain** | | 12.8 ± 0.6 | 49.9 ± 4.9** | 14.0 ± 1.5 | 46.6 ± 6.1^^^ | P<0.0001 | $F_{1,32}$ = 34.85 |
| **Hippocampus** | | 14.7 ± 3.1 | 75.8 ± 3.1*** | 13.7 ± 2.4 | 70.3 ± 14.2^^^ | P<0.0001 | $F_{1,37}$ = 45.05 |
| **Cortex** | | 15.7 ± 1.0 | 65.0 ± 5.7**** | 17.4 ± 1.7 | 61.4 ± 10.4^^^^ | P<0.0001 | $F_{1,43}$ = 57.73 |
| **Frontal Cortex** | | 26.3 ± 1.6 | 94.1 ± 5.5*** | 24.7 ± 2.7 | 96.5 ± 20.3^^^ | P<0.0001 | $F_{1,42}$ = 39.76 |

**Kynurenine-induced elevation in kynurenine levels in plasma and across brain regions.**
All data are mean ± SEM. Two-way ANOVA, Bonferroni's post hoc analysis:
Vehicle vs. Kynurenine: **P < 0.01, ***P < 0.001, ****P < 0.0001
PF-04859989 vs. PF-04859989 + Kynurenine: ^P < 0.05, ^^P < 0.01, ^^^P < 0.001, ^^^^P < 0.0001.

**Supplemental Table 2. Kynurenine challenge significantly increases kynurenine levels in plasma and across brain regions.** Adult rats were peripherally injected at Zeitgeber time (ZT) 0 with kynurenine (100 mg/kg) to induce de novo kynurenic acid (KYNA) formation. PF-04859989 (30 mg/kg), systemically active KAT II inhibitor, was given 30 minutes prior at ZT 23.5. Tissues were harvested at ZT 2. Kynurenine levels were evaluated. All data are mean ± SEM. N = 4 - 12 per group.

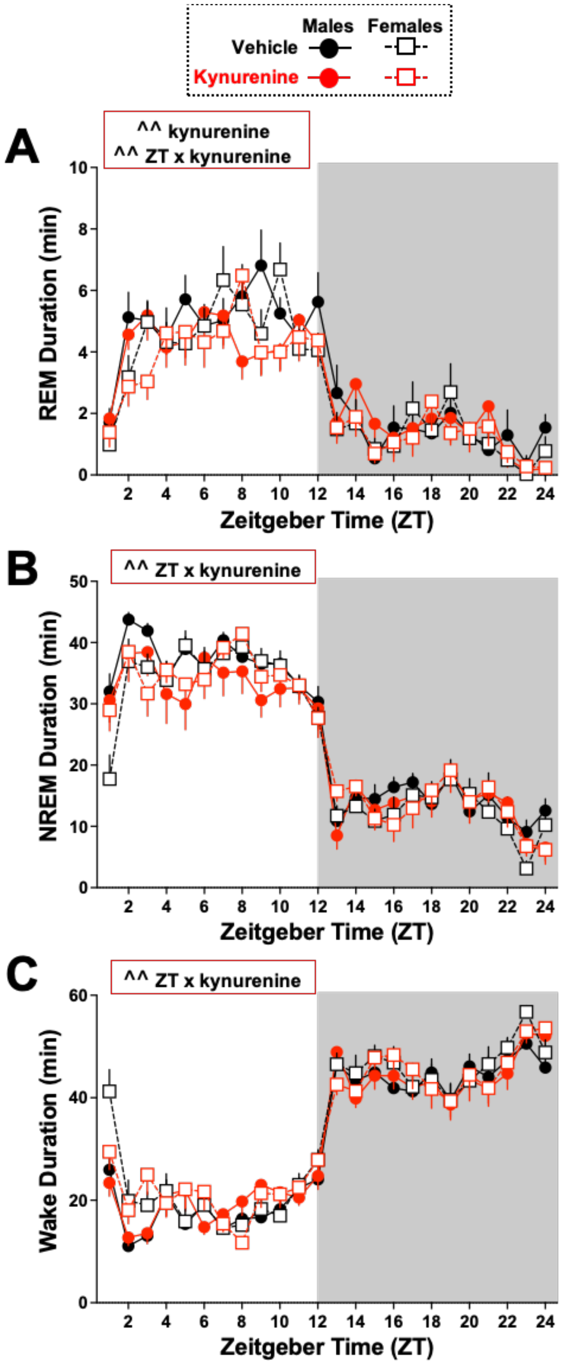

**Supplemental Figure 1. Impacts of kynurenine challenge on vigilance state duration.** Adult rats were treated with vehicle or kynurenine (100 mg/kg) at Zeitgeber time (ZT) 0. Data are mean ± SEM, analyzed by three-way RM ANOVA, with significance shown in red boxes in graphs as ^^P<0.01. N= 8 males, 12 females.

**(A)** 1-hr bins of REMS duration. ZT x Treatment: $F_{23,414}$ = 1.618, P<0.01; Treatment: $F_{1,18}$ = 15.44, P<0.01; ZT: $F_{23,414}$ = 29.37, P<0.0001. Sex: $F_{1,18}$ = 0.4279, P=0.5213.

**(B)** 1-hr bins of NREMS duration. ZT x Treatment: $F_{23,414}$ = 1.789, P<0.01 ZT: $F_{23,414}$ = 66.12, P<0.0001. Sex: $F_{1,18}$ = 0.3687, P=0.5513.

**(C)** 1-hr bins of wake duration. ZT x Treatment: $F_{23,414}$ = 1.807, P<0.01; ZT: $F_{23,414}$ = 70.51, P<0.0001. Sex: $F_{1,18}$ = 2.327, P=0.1445.

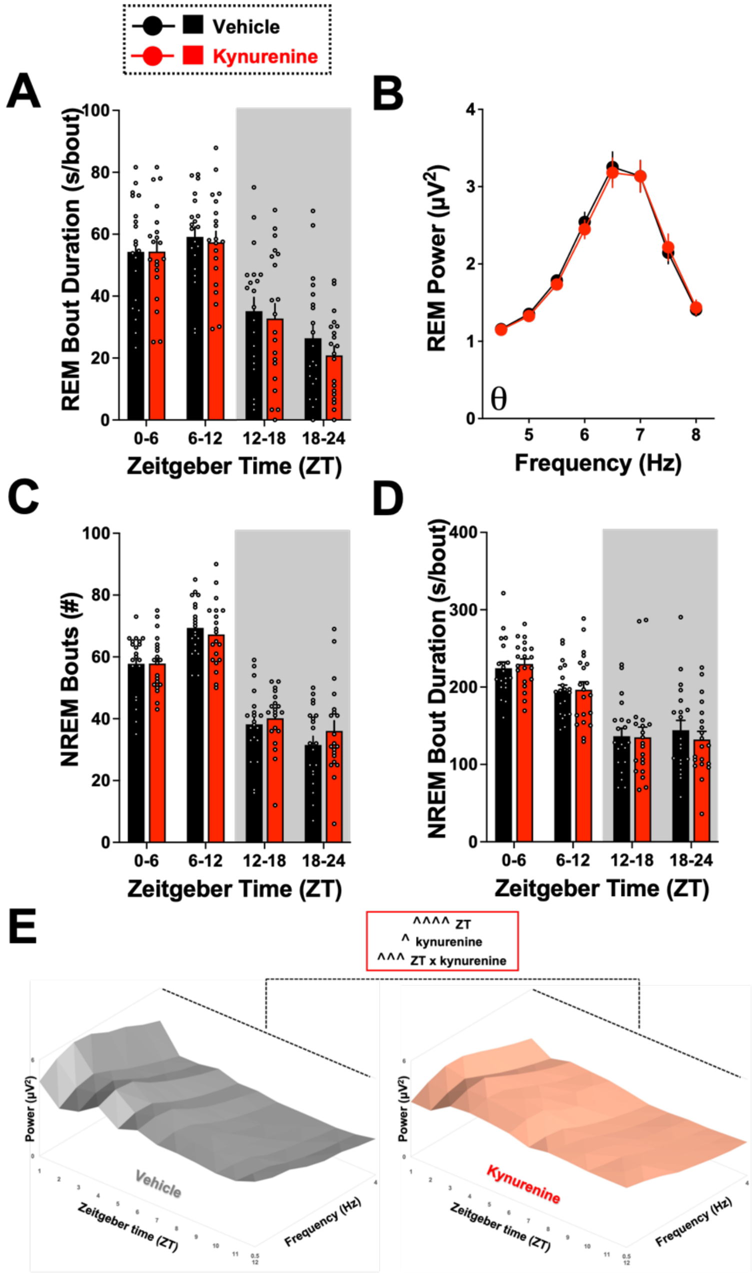

**Supplemental Figure 2. Impacts of kynurenine challenge on sleep architecture parameters.** Adult rats were treated with vehicle or kynurenine (100 mg/kg) at Zeitgeber time (ZT) 0. Data are mean ± SEM, analyzed by two-way RM ANOVA. **(A)** 6-hr bins of average REMS bout duration. ZT: $F_{3,57}$ = 48.65, P<0.0001. **(B)** REMS spectral power in the theta frequency range during the light phase. Frequency: $F_{7,133}$ = 54.11, P<0.0001. **(C)** 6-hr bins of NREMS bout number. ZT: $F_{3,57}$ = 75.97, P<0.0001. **(D)** 6-hr bins of average NREMS bout duration. ZT: $F_{3,57}$ = 44.36, P<0.0001. **(E)** Delta spectra power (0.5 – 4 Hz) normalized to total spectra power visualized in 1-hr bins. ZT: $F_{11,187}$ = 15.98,

P<0.0001. Treatment: $F_{1,17}$ = 4.615, P<0.05. ZT x treatment: $F_{11,187}$ = 3.233, P<0.001. N = 20 per group (8 males, 12 females).

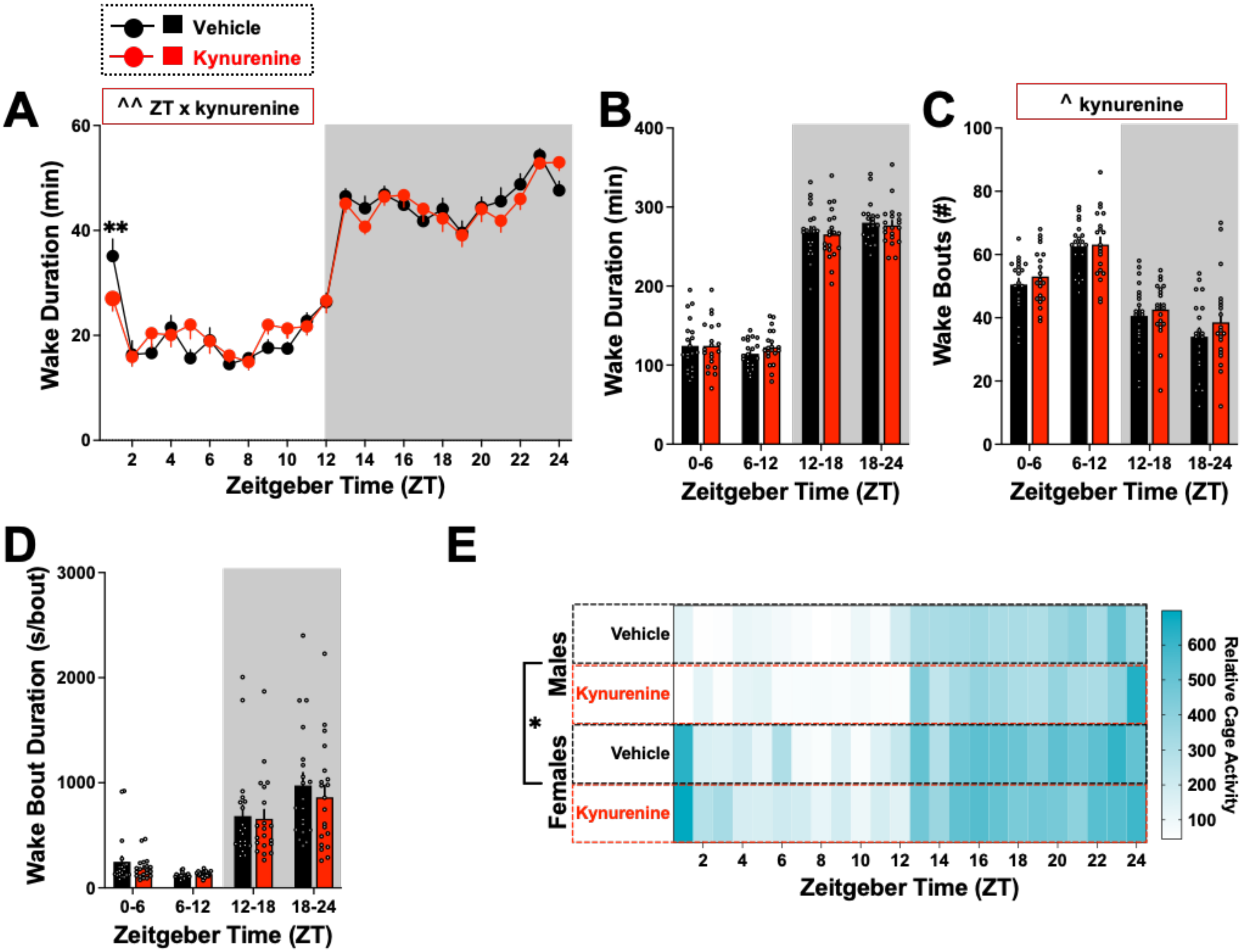

**Supplemental Figure 3. Impacts of kynurenine challenge on wake parameters.** Adult rats were treated with vehicle or kynurenine (100 mg/kg) at Zeitgeber time (ZT) 0. Data are mean ± SEM, analyzed by two-way RM ANOVA (A-D), with significance shown in graphs as ^P<0.05, ^^P<0.01 and Bonferroni's post hoc as **P<0.01 or three-way RM ANOVA (E), with significance shown in graph as *P<0.05. **(A)** 1-hr bins of wake duration. ZT x treatment: $F_{23,437}$ = 2.078, P<0.01. ZT: $F_{23,437}$ = 72.67, P<0.0001. **(B)** 6-hr bins of wake duration. ZT: $F_{3,57}$ = 265.5, P<0.0001. **(C)** 6-hr bins of wake bout number. Treatment: $F_{1,19}$ = 4.967, P<0.05. ZT: $F_{3,57}$ = 44.76, P<0.0001. **(D)** 6-hr bins of average duration per wake bout. ZT: $F_{3,57}$ = 34.07, P<0.0001. **(E)** Relative home cage activity. Sex: $F_{1,347}$ = 5.118, *P<0.05. ZT: $F_{22,396}$ = 25.40, P<0.0001. N = 20 per group (8 males, 12 females).

| Transition | Vehicle | | Kynurenine | | |
|---|---|---|---|---|---|
| **Light Phase** | | | | | |
| **Wake-NREMS** | 109 | ± 3.1 | 112 | ± 3.6 | |
| **REMS-Wake** | 46 | ± 1.4 | 46 | ± 1.3 | |
| **REMS-NREMS** | 11 | ± 1.1 | 7 | ± 0.9 | **** |
| **NREMS-Wake** | 63 | ± 2.5 | 66 | ± 2.9 | |
| **NREMS-REMS** | 57 | ± 1.6 | 53 | ± 1.4 | ** |
| **Dark Phase** | | | | | |
| **Wake-NREMS** | 66 | ± 4.6 | 77 | ± 3.8 | * |
| **REMS-Wake** | 15 | ± 1.7 | 18 | ± 2.2 | * |
| **REMS-NREMS** | 3 | ± 1.0 | 2 | ± 0.5 | |
| **NREMS-Wake** | 51 | ± 4.7 | 59 | ± 4.8 | |
| **NREMS-REMS** | 17 | ± 2.3 | 20 | ± 2.4 | |

Paired Student's t-test
* P<0.05, ** P<0.01, **** P<0.0001

**Supplemental Table 3. Acute kynurenine challenge disrupts female sleep transitions.** Adult male and female rats were treated with vehicle or kynurenine (100 mg/kg) at Zeitgeber time (ZT) 0. All data are mean ± SEM, analyzed by paired Student's t-test. N = 20 per group (8 males, 12 females).

**Supplemental Figure 4. Kynurenine challenge significantly influenced NREM sleep and REM sleep bout length distribution.** Adult rats were treated with vehicle or kynurenine (100 mg/kg) at Zeitgeber time (ZT) 0. Data are mean ± SEM, analyzed by two-way RM ANOVA (A-c), with significance shown in graphs as ^^P<0.01, ^^^^P<0.0001 and Bonferroni's post hoc as *P<0.05, **P<0.01. **(A)** Time bins of wake bouts. Time bin: $F_{6,66}$= 105.7, P<0.0001. **(B)** Time bins of NREM sleep bouts. Time bin: $F_{6,66}$= 112.6, P<0.0001. Kynurenine: $F_{1,11}$= 10.17, P<0.01. **(C)** Time bins of REM sleep bouts. Time bin: $F_{6,66}$= 34.60, P<0.0001. Kynurenine: $F_{1,11}$= 11.01, P<0.01. N = 12 per group (5 males, 7 females).

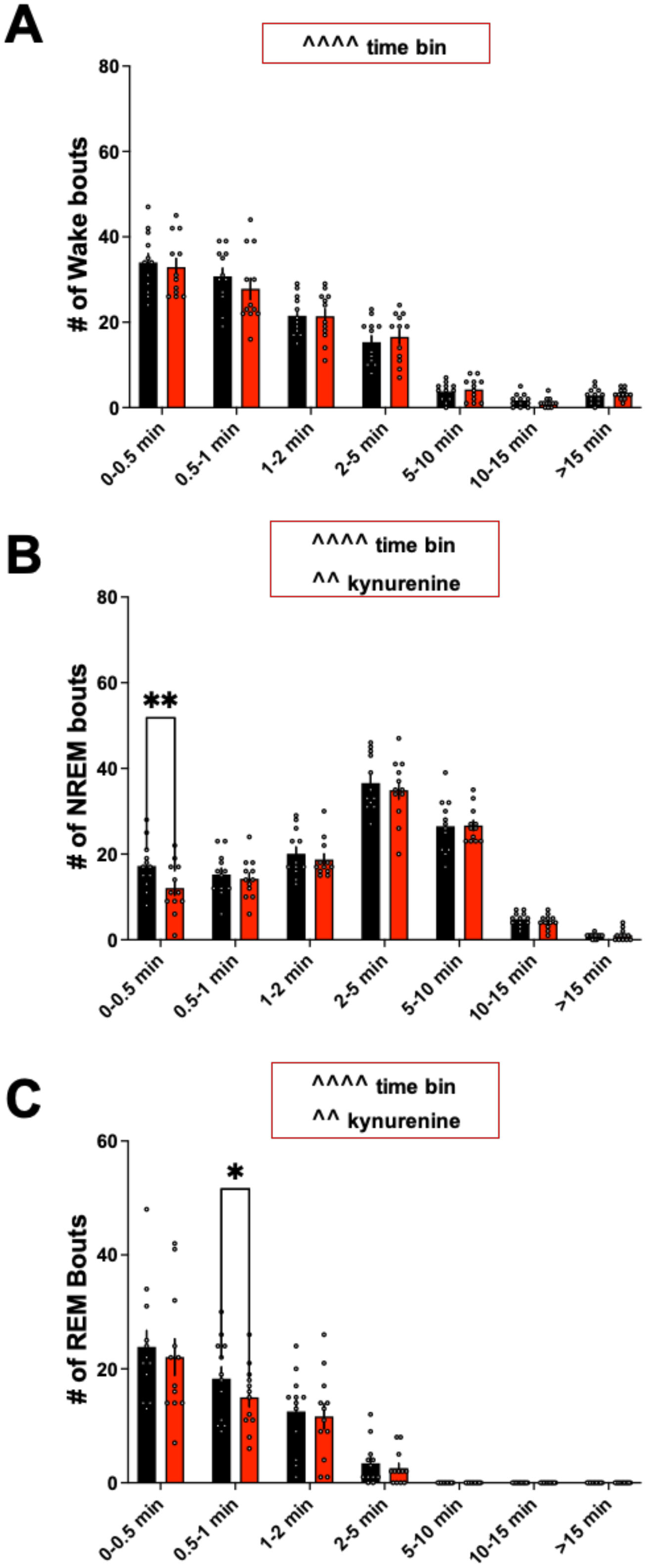

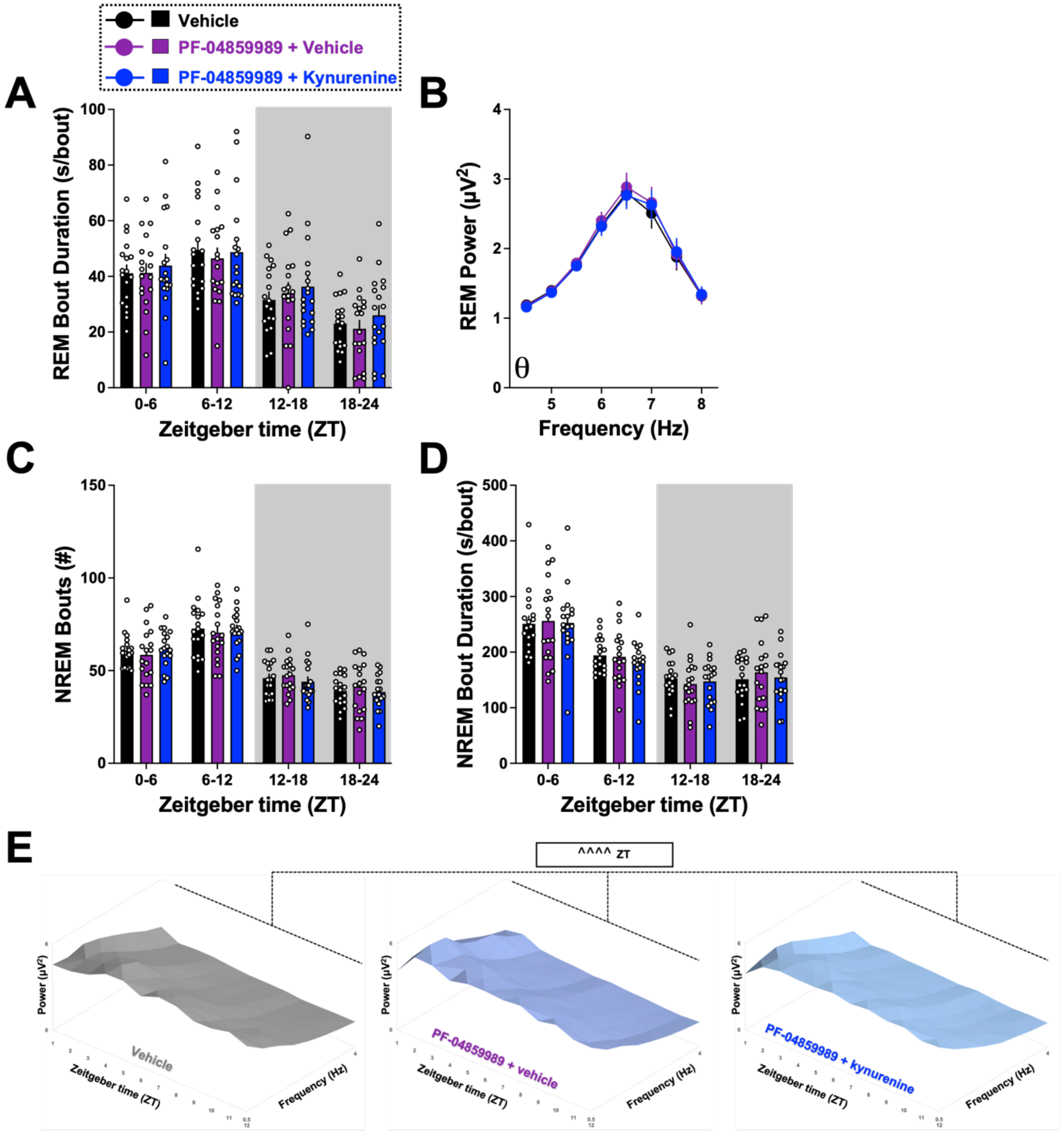

**Supplemental Figure 5. Impacts of PF-04859989 treatment prior to kynurenine challenge on sleep architecture parameters.** Adult rats were treated with vehicle, PF-04859989 (30 mg/kg), or PF-04859989 + kynurenine (100 mg/kg) at the start of the light phase. Data are mean ± SEM, analyzed by RM ANOVA. **(A)** 6-hr bins of average REMS bout duration. ZT: $F_{3,51}$ = 36.20, P<0.0001. **(B)** REMS spectral power in the theta frequency range during the light phase. Frequency: $F_{7,119}$ = 29.01, P<0.0001. **(C)** 6-hr bins of NREMS bout number. ZT: $F_{3,51}$ = 62.97, P<0.0001. **(D)** 6-hr bins of average NREMS bout duration. ZT: $F_{3,51}$ = 68.22, P<0.0001. **(E)** Delta spectra power (0.5 – 4 Hz) normalized to total spectra power visualized in 1-hr bins. ZT: $F_{11,143}$ = 10.80, P<0.0001. N = 18 per group (9 males, 9 females).

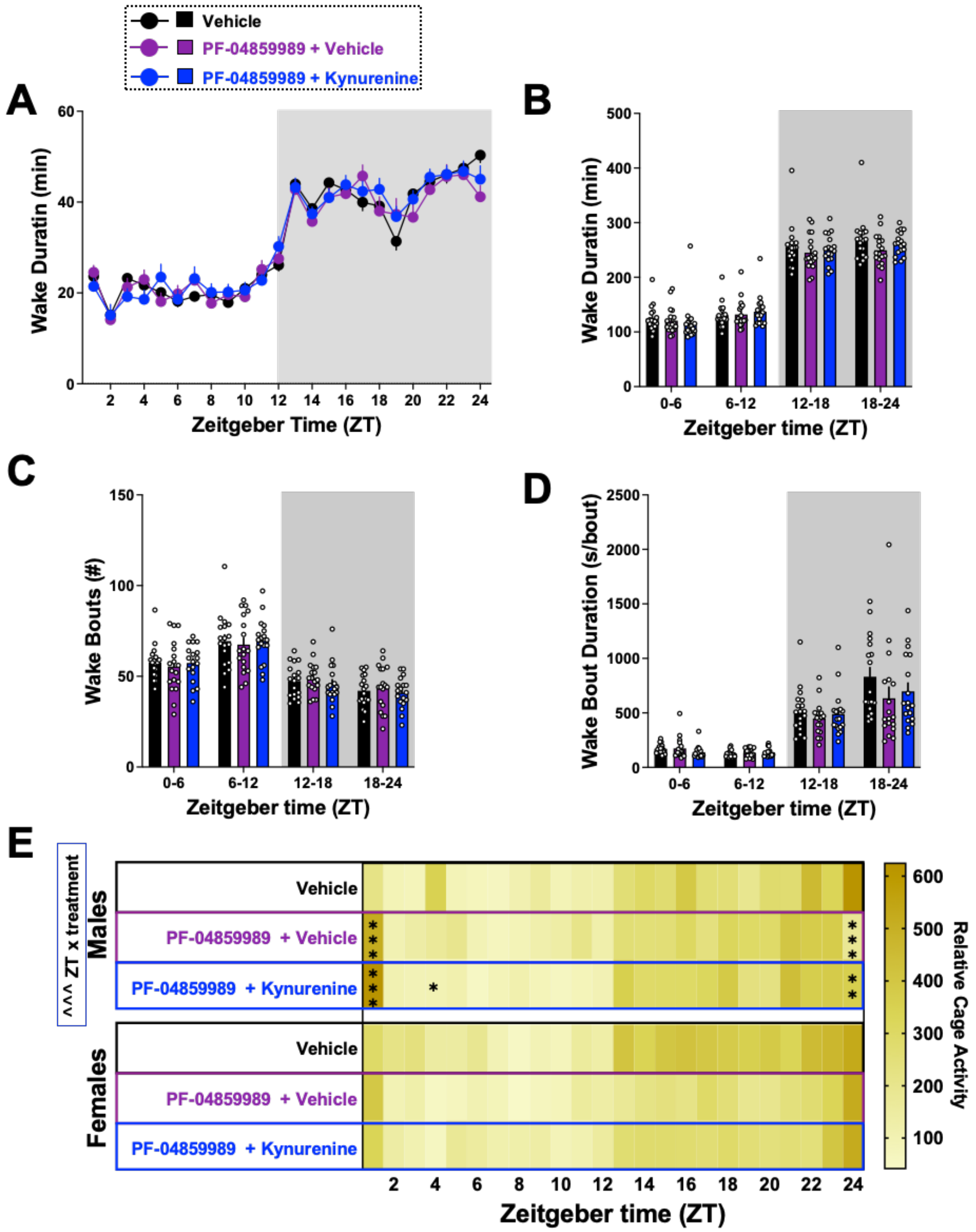

**Supplemental Figure 6. Impacts of pretreatment with PF-04859989 prior to kynurenine challenge on wake parameters.** Adult rats were treated with vehicle, PF-04859989 (30 mg/kg), or PF-04859989 + kynurenine (100 mg/kg) at the start of the light phase. Data are mean ± SEM, analyzed by ANOVA, with significance shown in graphs as ^^^P<0.001, and Dunnett's post hoc test *P<0.05, **P<0.01, ***P<0.001 vs vehicle. **(A)** 1-hr bins of wake duration. ZT: $F_{23,391}$ = 87.96, P<0.0001. **(B)** 6-hr bins of wake duration. ZT: $F_{3,51}$ = 283.1, P<0.0001. **(C)** 6-hr bins of wake bout number. ZT: $F_{3,51}$ = 46.80, P<0.001. **(D)** 6-hr bins of average duration per wake bout. ZT: $F_{3,51}$ = 51.67, P<0.0001. **(E)** Relative home cage activity, evaluated separated by sex. Males, ZT x treatment: $F_{46,322}$ = 2.075, P<0.001; ZT: $F_{23,184}$= 11.66, P<0.0001. Females, ZT: $F_{23,513}$ = 13.83, P<0.0001. N = 18 per group (9 males, 9 females).

| Transition | Vehicle | | PF-04859989 + Vehicle | | PF-04859989 + Kynurenine | |
|---|---|---|---|---|---|---|
| **Light Phase** | | | | | | |
| **Wake-NREMS** | 118 | ± 4.1 | 117 | ± 6.5 | 122 | ± 4.8 |
| **REMS-Wake** | 37 | ± 1.9 | 39 | ± 2.9 | 37 | ± 2.0 |
| **REMS-NREMS** | 4 | ± 0.8 | 3 | ± 0.6 | 4 | ± 0.8 |
| **NREMS-Wake** | 81 | ± 3.3 | 79 | ± 6.1 | 85 | ± 5.1 |
| **NREMS-REMS** | 41 | ± 1.9 | 42 | ± 3.1 | 41 | ± 2.1 |
| **Dark Phase** | | | | | | |
| **Wake-NREMS** | 77 | ± 3.0 | 83 | ± 4.4 | 78 | ± 3.7 |
| **REMS-Wake** | 16 | ± 1.4 | 19 | ± 2.5 | 17 | ± 1.5 |
| **REMS-NREMS** | 1 | ± 0.2 | 1 | ± 0.2 | 1 | ± 0.3 |
| **NREMS-Wake** | 61 | ± 2.9 | 65 | ± 5.0 | 61 | ± 3.3 |
| **NREMS-REMS** | 17 | ± 1.4 | 20 | ± 2.6 | 18 | ± 1.6 |

**Supplemental Table 4. PF-04859989 treatment does not alter vigilance state transitions.** Adult rats were treated with vehicle, PF-04859989 (30 mg/kg), or PF-04859989 + kynurenine (100 mg/kg) at the start of the light phase. All data are mean ± SEM, analyzed by ANOVA. N = 18 per group (9 males, 9 females).

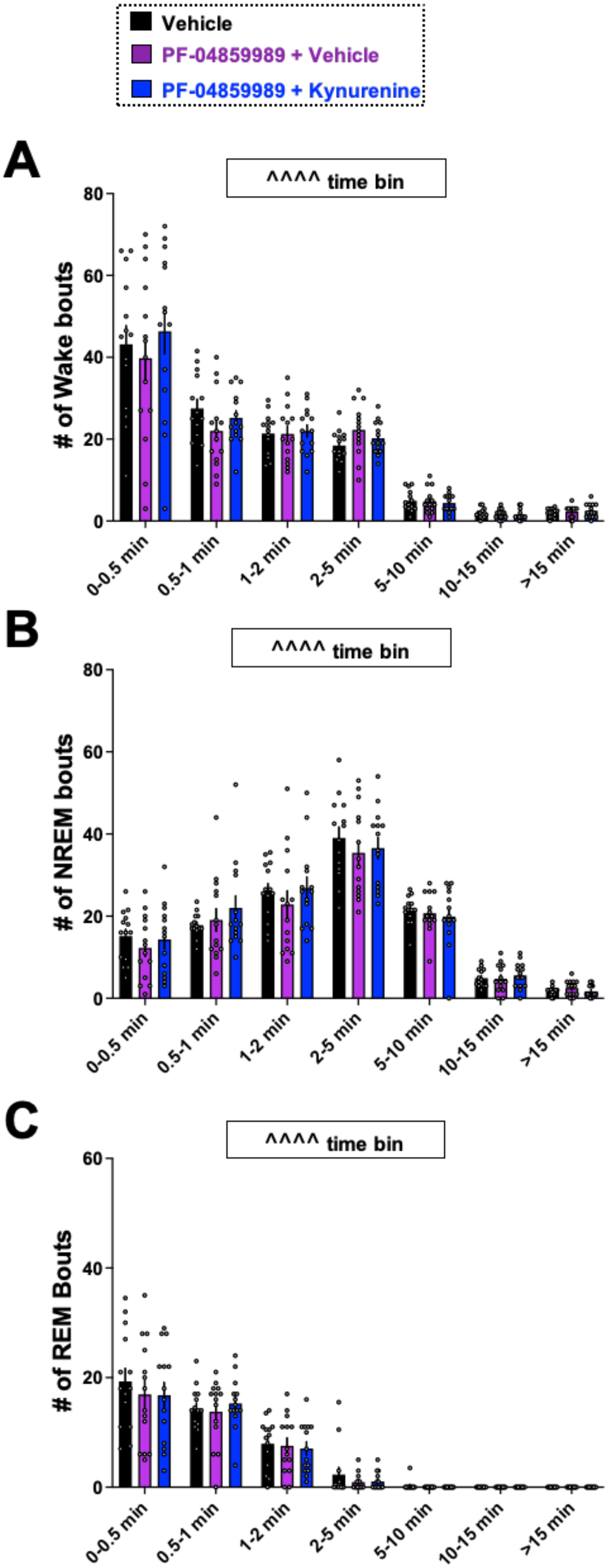

**Supplemental Figure 7. Impacts of pretreatment with PF-04859989 prior to kynurenine challenge on NREM sleep and REM sleep bout length distribution.** Adult rats were treated with vehicle, PF-04859989 (30 mg/kg), or PF-04859989 + kynurenine (100 mg/kg) at the start of the light phase. Data are mean ± SEM, analyzed by ANOVA. **(A)** Time bins of wake bouts. Time bin: $F_{6,78}$= 61.06, P<0.0001. **(B)** Time bins of NREM sleep bouts. Time bin: $F_{6,78}$= 79.41, P<0.0001. **(C)** Time bins of REM sleep bouts. Time bin: $F_{6,78}$= 55.26, P<0.0001. N = 14 per group (7 males, 7 females).
